# *In vitro* model of human subcutaneous adipocyte spheroids for studying mitochondrial dysfunction and mitochondria activating compounds

**DOI:** 10.1101/2025.04.16.649074

**Authors:** A Wagner, JH Lautaoja-Kivipelto, K Pehkonen, A Hassinen, M Kuusela, L Röttger, E Herbers, A Ioannidou, S Mädler, I Rothenaigner, S Srinivasan, S Laasonen, MT Rahman, P Elomaa, S Kortetjärvi, A Olsson, O Ukkola, K Hadian, M Mann, H Peltoniemi, KH Pietiläinen, M Klingenspor, KA Virtanen, CE Hagberg, E Pirinen

**Author notes:** Correspondence to Eija Pirinen, Research Unit of Biomedicine and Internal Medicine, Faculty of Medicine, University of Oulu, Oulu, Finland.

## Abstract

Mitochondrial abnormalities drive subcutaneous white adipose tissue dysfunction in obesity, leading to metabolic complications. A key challenge in obesity research is the lack of *in vitro* models to study human adipocyte mitochondria. Here, we establish a human subcutaneous adipocyte spheroid model to characterize mitochondrial metabolism under obesity-relevant conditions and drug exposure.

Human preadipocyte spheroids were differentiated in ultra-low attachment plates for 3 weeks in media free of thiazolidinediones. The differentiated adipocyte spheroids showed increased lipid accumulation, adipogenic gene expression, mitochondrial respiration and adiponectin secretion, and a proper response to hormonal stimulation. Lipid mixture administration during differentiation induced metabolic disturbances including mitochondrial respiration failure associated with increased mitochondrial biogenesis. Post-differentiation treatment with rosiglitazone, a peroxisome proliferator-activated receptor γ agonist, improved mitochondrial bioenergetics and adiponectin secretion in lipid mixture-administered adipocyte spheroids.

Our model enables precise measurement of adipocyte mitochondria under various conditions, providing a platform for mitochondria-related research and drug discovery in obesity.

## INTRODUCTION

Subcutaneous white adipose tissue (scWAT) represents the major triglyceride store of the body. The metabolic and endocrine functions of scWAT play a crucial role in regulating energy balance and controlling systemic lipid and glucose homeostasis (1). Fatty acids are mobilized from lipid droplets by lipolysis to fuel the energy demand in other tissues, while adipokines secreted by adipocytes improve fatty acid and glucose utilization in target tissues (1). In obesity, scWAT functions are disturbed when adipocyte triglyceride storage exceeds expansion capacity, leading to ectopic lipid accumulation and subsequently to the development of metabolic complications, such as insulin resistance in non-adipose tissues (2). Thus, treatments restoring the key metabolic and endocrine functions in scWAT could help to reconstitute metabolic health in obesity.

One of the key factors triggering scWAT dysfunction in obesity is mitochondrial abnormalities (3). Mitochondrial alterations, including reduced mitochondrial respiration, number and mass, as well as distrupted dynamics, have been reported in scWAT of humans and rodents (3, 4). ScWAT mitochondrial dysfunction has been proposed to be a consequence of chronic energy surplus-induced inflammation (3). However, the understanding of the pathological causes leading to the development of scWAT mitochondrial dysfunction have remained incomplete. As scWAT mitochondrial dysfunction occurs in the early-stage of weight gain and obesity (6), even before the onset of obesity-related metabolic complications (8), it is plausible that the malfunction of scWAT mitochondria contributes to the deterioration of the whole-body glucose and lipid homeostasis. In support, the activation of tissue mitochondrial metabolism protects against obesity-related metabolic complications in mice (5). Clinically, pharmacological targeting of mitochondria is still an unmet need with great potential health benefits. Overall, the identification of the molecular mechanisms resulting in scWAT mitochondrial dysfunction as well as mitochondria activating compounds could markedly advance the drug development for obesity-related metabolic complications.

One of the main limitations in obesity research and the drug discovery field has been the lack of physiological *in vitro* models for human white subcutaneous adipocytes. The current *in vitro* research on human subcutaneous adipocytes mostly relies on differentiation of precursor cells and preadipocytes into lipid-laiden adipocytes using two-dimensional (2D) monolayer cultures (7). However, human precursor cells and preadipocytes fail to differentiate and mature efficiently into white adipocytes with an unilocular lipid droplet morphology and metabolic function in these monolayer cultures. For example, differentiated adipocytes can acquire brown adipocyte-like properties, such as multilocular lipid droplet morphology, high thermogenic activity and/or uncoupling protein 1 (UCP1) expression (9, 10). In monolayer cultures, these brown adipocyte-like properties can be caused by the use of high concentration of thiazolidinediones, such as peroxisome proliferator-activated receptor γ (PPARγ) agonist rosiglitazone, which promote adipocyte differentiation. By solving some of these technical challenges, three-dimensional (3D) culture models hold great promise over traditional 2D monolayer cultures (9, 11). Several recently developed human white adipocyte spheroid models (9, 12) report 3D cultured adipocytes to better resemble their *in vivo* counterparts, showing more unilocular lipid droplet morphology and higher expression of adipogenic genes and adipokine secretion in comparison to 2D monolayer cultures (11). However, none of the previously developed 3D models of human white adipocytes have been optimized to specifically study mitochondrial (dys)function and testing of mitochondria targeting pharmacological compounds.

The objective of this study was to establish a human subcutaneous white adipocyte spheroid culture model that allows the investigation of mitochondrial variables under various metabolic conditions and in response to mitochondrial-activating drugs. We demonstrate that our model effectively mimics the mitochondrial remodeling associated with adipogenic differentiation and lipid overloading-induced mitochondrial dysfunction. This enables the assessment of human white subcutaneous adipocyte mitochondrial biology in both physiological and pathophysiological contexts, respectively. Moreover, we present our adipocyte spheroid model as a valuable *in vitro* platform for investigation of pharmacological compounds that may influence mitochondrial function, thereby advancing novel drug discovery and deepening our understanding of disease mechanisms.

## METHODS

### Tissue and cell line donor characteristics

Human preadipocytes isolated from scWAT from two healthy (e.g. no diabetes) female Caucasian donors with body mass index (BMI) 23 (Cat No. SAT-PT-5020, TAN 33226) and 28 (Cat No. PT-5020, TAN 27458) were purchased from Lonza (Basel, Switzerland) (Table 1, donors 1 and 2). The preadipocytes from both donors were tested negative against mycoplasma using EZ-PCR Mycoplasma Detection Kit (20-700-20, Biological Industries, Kibbutz Beit-Haemek, Israel).

**TABLE 1.** The donor characteristics for subcutaneous preadipocytes, mature adipocyte fraction (MAF) cells and subcutaneous white adipose tissue (scWAT) biopsies. BMI, body mass index. Age is reported at the time of the sample collection.

|  | **Cell/tissue type donated** | **Sex** | **Age (years)** | **Weight (kg)** | **Height (cm)** | **BMI** |
| --- | --- | --- | --- | --- | --- | --- |
| Donor 1 | Preadipocytes | Female | 43 | 65 | 168 | 23 |
| Donor 2 | Preadipocytes | Female | 42 | 77 | 165 | 28 |
| Donor 3 | scWAT/MAF | Female | 43 | 66 | 161 | 26 |
| Donor 4 | MAF | Female | 52 | 68 | 165 | 25 |
| Donor 5 | MAF | Female | 49 | 90 | 174 | 30 |
| Donor 6 | scWAT | Female | 40 | 81 | 170 | 28 |
| Donor 7 | scWAT/MAF | Female | 36 | 53 | 168 | 19 |
| Donor 8 | scWAT/MAF | Female | 53 | 57 | 156 | 23 |
| Donor 9 | scWAT/MAF | Female | 51 | 59 | 156 | 24 |

Human mature adipocyte fraction cells (MAFs) and scWAT samples were obtained from scWAT biopsies collected from the abdominal area of seven healthy (e.g., no diabetes) female Caucasian donors who underwent an elective liposuction operation at Tilkka Hospital, Helsinki (Table 1, donors 3 to 9). Immediately after surgery, scWAT was dissected to remove connective tissue and blood vessels. One part of the scWAT was snap-frozen in liquid nitrogen and stored at -80°C until use, while another part was used for MAF cell harvesting. Isolation of MAFs was performed using the previously described mechanical and enzymatic method (13). All the donors provided written informed consent, and the study protocol was approved by the Ethics Committee of Helsinki University Hospital (HUS/1039/2019).

### Cell culture

Subcutaneous preadipocytes were cultured in growth media (3.15 g/L glucose, 21331, Gibco, Thermo Fisher Scientific, Waltham, MA, USA) supplemented with 5% human serum (S4190, Biowest, Nuaillé, France), 1% glutamax (35050, Gibco, Thermo Fisher Scientific), 15 ng/mL amphotericin B (A2924, Sigma-Aldrich, Saint Louis, MO, USA) and 30 μg/mL gentamicin (G1397, Sigma-Aldrich) in a humidified environment at 37°C and 5% CO_2_. Preadipocytes were expanded typically 4 to 6 cycles before seeding for the experiments. After removal of serum-containing growth media by washing the cells with 1x Dulbecco’s phosphate buffered saline (DPBS) or 1x phosphate buffered saline (PBS), adipocyte spheroids were created by seeding 15 000, 25 000 or 35 000 preadipocytes in 200 µL per well in serum-free growth media in a 96-well plate with a cell repellent surface (650970, Greiner Bio-One GmbH, Kremsmünster, Austria). The spontaneously formed spheroids collected after 2 days served as undifferentiated preadipocyte spheroids. For the generation of differentiated adipocyte spheroids, 150 µL of the growth media was removed and differentiation was induced by adding 150 µL of serum- and PPARγ agonist thiazolidinedione-free adipocyte differentiation media (DM, 1 g/L glucose, 1 µmol dexamethasone, 1 µg/mL human insulin, 500 µmol 3-isobutyl-1-methylxanthine and 100 µmol indomethacin and other compounds from the media kit (14, 15) PB-C-DH-442-3699, PELOBiotech GmbH, Planegg, Germany). Alternatively, after removal of growth media, adipocyte spheroids were embedded in 30 µL of growth factor reduced (GFR)-Matrigel (356231, Corning Inc. Corning, NY, USA) followed by a 30-min incubation at 37°C and 5% CO_2_ before administration of 150 µL of DM per well. Independent of the differentiation initiation, media was partly (3/4) changed every two to three days until day 7 (2-week protocol), day 14 (3-week protocol) or day 20 (4-week protocol). Next, DM was replaced by adipocyte maintenance media (MM, 1 g/L glucose, 1 µg/mL insulin, PB-C-MH-442-3699-M, PELOBiotech GmbH) which was partly (3/4) refreshed every two to three days until day 14 (2-week protocol), day 21 (3-week protocol) or day 28 (4-week protocol). Notably, instead of using the antibiotics provided by the DM and MM kits, DM and MM media were supplemented with amphotericin B and gentamycin similarly as described for the growth media. The 3-week protocol with 35 000 preadipocytes per well embedded in GFR-Matrigel was selected as the gold standard protocol and this was used throughout this article if not stated otherwise.

To mimic nutrient-based energy surplus-induced fat accumulation and metabolic complications, adipocyte spheroids from the donor 2 with BMI 28 were incubated with 1:125 chemically defined lipid concentrate (*i.e*., lipid mixture (LM), 11905031, Thermo Fisher Scientific) throughout the 3-week protocol including both differentiation and maintenance phases. Furthermore, to test the suitability of our model for pharmacological studies, adipocyte spheroids were administered with DM and MM as described above or with DM and MM containing LM throughout the regular 3 week protocol, and thereafter the LM-free adipocyte spheroids were treated with vehicle (0.1% DMSO) for 72 hours, while the LM-administered adipocyte spheroids were treated with vehicle (0.1% DMSO) or 1 μM rosiglitazone in 0.1% DMSO (557366, Calbiochem, Sigma-Aldrich) for 72 hours.

### RNA extraction, cDNA synthesis and quantitative real-time PCR

For the RNA extraction, 12 to 15 adipocyte spheroids were pooled per one sample. In addition, RNA was isolated from MAFs and scWAT pieces from donors 3-9 (Table 1). The RNA from all the samples was extracted using TRIzol™ Reagent (15596018, Invitrogen, Waltham, MA, USA). The adipocyte spheroids were homogenized using the Kimble® Pellet Pestle® Cordless Motor (DWK Life Sciences GmbH, Germany) with appropriate tissue grinders (DWK Life Sciences GmbH, Germany), while scWAT and MAF samples were homogenized using TissueLyser LT (4 min, 45 Hz, Qiagen). Thereafter, all the samples were centrifuged for 10 min at 12 000 x g at 4°C. Next, chloroform (20% of the initial sample volume) was added to the supernatant, samples were mixed and incubated at room temperature (RT) for 3 min before centrifugation for 30 min at 12 000 x g at 4°C. The upper organic phase was transferred into a new tube and equal volume of 80% ethanol was added before gentle mixing. From this step onwards, the protocol of RNeasy Micro Kit (74004, Qiagen, Hilden, Germany) for RNA extraction from adipocyte spheroids or the protocol of RNeasy Mini Kit (74104, Qiagen) for RNA extraction from scWAT and MAF was followed according to the manufacturer’s protocol without the use of DNase. The reverse transcription was conducted to all the samples using SuperScript™ VILO™ cDNA Synthesis Kit (10499763, Invitrogen) according to the manufacturer’s instructions. The quantitative real-time polymerase chain reaction (RT-qPCR) was conducted according to the manufacturer’s protocol with SYBR Green Master Mix (K0252, Thermo Fisher Scientific). The template amount used for RT-qPCR was 15 ng per well, independent of the sample type. The CFX96 Real-Time PCR Detection System combined with CFX Maestro software (Bio-Rad Laboratories Inc., Hercules, CA, USA) as well as efficiency corrected ΔΔCt method were used for data analysis. Of the tested housekeeping genes, importin 8 (*IPO8*) was selected for the data normalization due to the unaltered expression and the lowest variation among the groups. The primers used in the experiments are listed in Table S1.

### Fluorescence live cell imaging

Adipocyte spheroids were imaged using thePerkinElmer Opera Phenix® High-Content Screening confocal system. Prior to imaging, adipocyte spheroids were washed twice with 1x PBS and incubated in a staining solution consisting of 1x DPBS with 0.5 µg/mL BODIPY (D3922, Thermo Fisher Scientific) to label lipid droplets, 1 µg/mL Hoechst (H3570, Sigma-Aldrich) to label nuclei and 100 nM MitoTracker Red CMXRos (M7512, Thermo Fisher Scientific) to label mitochondria. The samples were placed on a shaker at 30 rpm for 20 min and protected from light. Next, four spacers (IS007, iSpacer 0.2mm, SunJin Lab, Hsinchu City, Taiwan) were placed in the corners of a glass slide and 100 µL of Aguatex® (108562, Merck) was added in the middle. Adipocyte spheroids were washed twice with 1x DPBS for 5 min, placed into Aquatex®, and covered with a cover slip. The slides were then sealed with a non-fluorescent transparent nail polish. Afterwards, adipocyte spheroids were imaged using a 40x air objective (NA 0.6) or a 40x water immersion objective (NA 1.1) with a PerkinElmer Opera Phenix spinning disk confocal microscope. The following channels were used for the imaging: 350 for Hoechst, 488 for BODIPY and 568 for MitoTracker. To quantify the fluorescence intensity of the lipid droplets, mitochondria, and nuclei in adipocyte spheroids, confocal images were analyzed using Harmony 4.9 software (PerkinElmer) with a pipeline specified in Table S2. Briefly, the confocal images were analyzed to quantify the diameter, area and number of lipid droplets, and the number, mass and morphology of mitochondria within the (pre)adipocyte spheroids. Differentiation efficiency was calculated as the percentage of differentiated cells relative to the total cell number within each adipocyte spheroid (non-lipid bearing cells were not excluded beforehand). For the other lipid droplet parameters, the datasets were analyzed with a more robust background correction, particularly in the case of LM-administered adipocyte spheroids, compared to the analysis of mitochondrial parameters or differentiation efficiency. To remove false positive cells in the spheroids, a threshold was set based on the average mean lipid length (in µm) in both donor 1 and donor 2 undifferentiated spheroid datasets. The threshold was 2.27 µm for donor 1 and 0 µm for donor 2. All cells over the thresholds were removed from the undifferentiated datasets before proceeding to the data analysis. Only the cells in differentiated adipocyte spheroids containing lipids; with a value of over the threshold, were included in the differentiated spheroid dataset analysis of lipid droplet parameters. For the mitochondrial-related parameters, the cells with a value of 0 in the mitochondrial data were excluded, before a subsequent data analysis was performed.

### Mitochondrial respiration

Mitochondrial respiration measurements were performed using the Agilent Seahorse XFe96 Analyzer (Agilent Technologies, Santa Clara, CA, USA). First, the calibration cartridge (Agilent Seahorse XFe96/XF Pro Extracellular Flux Assay Kits, Agilent Technologies) was incubated with Agilent Seahorse XF Calibrant solution (100840-000, Agilent Technologies) overnight at 37°C in a humified environment the day before the run according to the manufacturer’s instructions. On the day of the run, adipocyte spheroids were collected, washed with 1x PBS, and placed on a precoated (0.1 mg/mL poly-D-glycine, P6407, Sigma-Aldrich) Agilent Seahorse XFe96 Spheroid Microplate (102978-100, Agilent Technologies). The media used in the runs was as follows: Seahorse base media (103335-100, Agilent Technologies) freshly supplemented with 10 mM glucose (10141520, Thermo Fisher Scientific), 2 mM glutamax (35050, Gibco) and 1 mM pyruvate (S8636, Sigma-Aldrich). All the experimental supplements, such as LM, DMSO, and/or rosiglitazone were added to the Seahorse base media with the identical concentrations as in the MM during the experiments. To ensure optimal conditions for adipocyte spheroids, Seahorse base media pH was verified to be 7.4 after addition of the supplements and other compounds. The Seahorse XF Cell Mito Stress Test Kit (103015-100, Agilent Technologies) was used according to the manufacturer’s instructions for the preparation of the injection compounds including oligomycin, an inhibitor of mitochondrial ATP synthase, carbonylcyanide-p-trifluoromethoxyphenylhydrazone (FCCP), a mitochondrial uncoupler, and rotenone and antimycin A, mitochondrial complex I and III inhibitors, respectively. The Seahorse run protocol was as follows; baseline 4-8 cycles, 2.5 µM oligomycin 10-18 cycles, 1.5 µM FCCP 12-18 cycles, and 1 µM rotenone and 1 µM antimycin A 12-25 cycles. Each cycle, consisting of a mixing, equilibrating and measuring step, lasted for 2 min. The number of cycles for each compound was optimized for each run separately to reach the oxygen consumption rate (OCR) plateau. After the run, the media were removed and adipocyte spheroids were washed with 1x PBS before storaging the samples at -80 °C until total DNA amount measurement for data normalization. The OCR of adipocyte spheroids was determined, and several variables of mitochondrial respiration including basal respiration, proton leak, maximal respiration, ATP-linked respiration, spare respiratory capacity and coupling efficiency were calculated as previously described by selecting the three highest consecutive OCR values for basal and maximal respiration, and the two to three lowest OCR values following inhibitor injection for each sample (16). The data was normalized using the Seahorse Wave Desktop software and finally processed in the Excel (Microsoft).

### Total DNA

For the measurement of total DNA, adipocyte spheroids were homogenized using a lysis buffer containing 0.5 M NaOH, 180 mM NaCl and 1 mM EDTA. After a 1- to 2-hour incubation at 37°C, samples were neutralized by 1 M Tris-HCl after which RNase A (EN0531, Thermo Fisher Scientific) was added to a final concentration of 100 µg/mL. Next, total DNA amount was measured using CyQUANT™ Cell Proliferation Assay (C7026, Thermo Fisher Scientific) according to the manufacturer’s instructions with the exceptions of not using standards provided in the kit and using the above described lysis buffer and 1 M Tris-HCl in 1:9 ratio as a diluent for the CyQUANT™ GR detection dye. The samples were measured in black OptiPlate 96-well plates (6005270, PerkinElmer) with Tecan infinity M200 (Tecan Trading AG, Switzerland) at fluorescence settings of 480/520 nm (mitochondrial respiration normalization) or with PerkinElmer VICTOR3 V 1420 Multilabel Counter (Perkin Elmer, Boston, MA, USA) at 485/535 nm (lipolysis normalization).

#### Bright-field microscopy

(Pre)adipocyte spheroids were imaged on day 2 (undifferentiated) or day 21 (differentiated) form U-bottom 96-well plates using Leica MC120 HD microscope (4X magnification, Leica Microsystems GmbH, Wetzler, Germany) together with Lilliput imaging system (7’’ HD Camera monitor, Zhangzhou, China). The diameter of the adipocyte spheroids was analyzed using Fiji/ImageJ software (version 1.54f on Windows, National Institutes of Health (NIH), Bethesda, MD, USA).

#### Cell viability

The adipocyte spheroid media lactate dehydrogenase (LDH) enzyme activity as a measure of cell viability was analyzed using LDH kit (981906, Thermo Fisher Scientific) and an automated Indiko plus analyzer (Thermo Fisher Scientific) according to the manufacturer’s instructions.

#### Intracellular triglycerides

Intracellular triglyceride content from adipocyte spheroids was evaluated by using AdipoRed™ Assay Reagent (PT-7009, Lonza) according to the manufacturer’s instructions. For the analysis, individual adipocyte spheroids were dissociated using Accumax (07921, StemCell Technologies) as previously described (17) with slight modifications. Briefly, media was removed, adipocyte spheroids were washed with 1x PBS and they were transferred onto a black OptiPlate 96-well plate in a 30 µL volume. Next, 70 μl of Accumax was added per adipocyte spheroid and the plate was then placed on a thermomixer (37°C, 1200 rpm, PHMP, Grants Instruments Ltd, Shepreth, UK). The 40-min dissociation protocol consisted of four 10 min cycles during which chemical dissociation was enhanced by mechanical dissociation achieved by pipetting Accumax up and down 10 times every 10 min. The intracellular triglyceride content was analyzed using AdipoRed staining measured at 485/570 nm and the results were normalized against Hoechst 33342 (H3570, Thermo Fisher Scientific) staining measured at 355/460 nm using VICTOR2 Wallac 1420 Multilabel Counter (Perkin Elmer, Boston, MA, USA).

#### Adiponectin and MCP-1 ELISA

The concentration of adiponectin (DRP300, Human Total Adiponectin/Acrp30 Immunoassay, RnD Systems, Minneapolis, MN, USA) and monocyte chemoattractant protein 1 (MCP-1, ab179886, Human MCP-1 ELISA Kit, Abcam) were analysed from the media using ELISA assays. The media were collected from the differentiated adipocyte spheroids on differentiation day 21, 3 days after the last media change, while the media from the undifferentiated preadipocyte spheroids were collected after 2 days of culture. Media samples were stored at -80°C until use. All the analyses were conducted according to the manufacturer’s protocols. In the assays, the sensitivity for adiponectin was 0.891 ng/mL and for MCP-1 1.26 pg/mL.

#### Protein extraction and capillary Western blot

For the analysis of insulin signaling, adipocyte spheroids were incubated in insulin-free DMEM (11880, 1 g/L glucose, Gibco) supplemented with 1% glutamax for 24 hours after three weeks of differentiation (day 21). The next day (day 22), adipocyte spheroids were administered with 1x PBS (vehicle) or 100 nM human insulin (I9278, Sigma-Aldrich) for 30 min in fresh DMEM supplemented with 1% glutamax followed by washing with 1x PBS before storing the samples at -80°C until use. 1x PBS was used to dilute insulin. If the adipocyte spheroids were administered with rosiglitazone and/or LM, these compounds were added to the media for the 30 min incubation with equal concentrations as for the other experiments. The proteins were extracted by adding 50 µL RIPA buffer (50 mM Tris-HCl (pH 7.6), 150 mM NaCl, 1 mM EDTA, 1% Triton X-100, 0.5% deoxycholic acid (97%), 0.1% SDS) supplemented with 3% Halt protease and phosphatase inhibitor cocktail (78442, Thermo Fisher Scientific) per one adipocyte spheroid. Each adipocyte spheroid was mechanically homogenized by 2 x 10 sec dissociation by Bel-Art ProCulture Cordless homogenizer (17455799, Thermo Fisher Scientific) and to enhance chemical dissociation, the lysates were incubated on ice for 20 min before centrifugation at 16 000 x g at 4°C for 20 min. The lysates were mixed with a fluorescent master mix (loading buffer) in a 4:1 ratio, denatured at 95°C for 5 min and stored at 4°C until (less than two weeks) capillary Western blot. The protein analysis was conducted by capillary Western blot and the JESS system as previously described with slight modifications (18). Notably, the protein content of the adipocyte spheroids was not measured before loading due to the GFR-Matrigel interference on protein extraction. More specifically, GFR-Matrigel is composed of *e.g.*, different collagen proteins and co-extraction of this material would overestimate the adipocyte spheroid protein content. Instead, equal volume of protein lysate from the undifferentiated and differentiated (pre)adipocyte spheroid samples was loaded per capillary. The 12-230 kDa capillary cartridge (SM-W004, ProteinSimple, Bio-Teche, Minneapolis, MN, USA) and chemiluminescence (anti-Rabbit Detection Module, DM-001, ProteinSimple, Bio-Teche) were used for protein detection according to the manufacturer’s instructions. The following primary antibodies with specific dilutions were used: p-Akt^Ser473^ (9271, 1:10, Cell Signaling Technology, Danvers, MA, USA), Akt, (9272, 1:50, Cell Signaling Technology) and Vinculin (ab129002,1:250, Abcam). Vinculin was used for data normalization. The data was analysed using the Compass for Simple Western software version 6.2.0 (ProteinSimple, Bio-Teche).

#### Lipolysis

Adipocyte spheroid lipolysis rate was evaluated by quantifying glycerol release to the media. The measurements under basal and isoproterenol-induced conditions were conducted similarly as previously described (19) with small modifications, such as exclusion of water bath incubation. Briefly, the cells were washed with 1x PBS and incubated for 3 h at +37 °C and 5% CO_2_ in 30 µL of Krebs-Ringer lipolysis buffer (136 mM NaCl, 1 mM NaH_2_PO_4_, 1 mM CaCl_2_, 4.7 mM KCl, 1 mM MgSO_4_, 2 mM glucose, 25 mM HEPES and 2% fatty acid-free BSA, pH 7.4) with water (vehicle, basal condition) or 10 µM isoproterenol (isoproterenol hydrochloride, I6504, Sigma-Aldrich, stimulated condition). Water was used as isoproterenol dissolvent. The glycerol content from the media was measured using Free Glycerol Reagent (F6428, Sigma-Aldrich) mixed with Amplex UltraRed Reagent (A36006, Thermo Fisher Scientific) at a 1:100 ratio. The reaction was incubated for 15 min at RT and the fluorescence was measured at 550/581 nm using VICTOR2 Wallac 1420 Multilabel Counter (Perkin Elmer). The data were normalized to the total DNA amount described in the total DNA measurement section.

#### Mitochondrial and genomic DNA

For the mitochondrial DNA (mtDNA) and genomic DNA (gDNA) extraction, 13 to 15 adipocyte spheroids were pooled per one sample. To compare the undifferentiated and differentiated (pre)adipocyte spheroids, pooled samples were homogenized in 500 µL of ice-cold 1x PBS using Precellys ceramic beads (Precellys lysing kit, Hard tissue grinder KT03961-1-001-2) with a frequency of 6600 rpm for 2x 30 sec, with the Precellys homogenizer (Bertin Instruments, cat. P000669-PR240- A). For the other comparisons, adipocyte spheroids were lysed using TissueLyser II (Qiagen) with variable times of 50 sec for the undifferentiated preadipocyte spheroids and 1.5 min for the differentiated adipocyte spheroids at a frequency of 20 Hz. The homogenization was followed by a 8 min centrifugation at 5000 x g at 4°C. The pellets were resuspended into a lysis buffer (10 mM Tris, 1 mM EDTA, 0.3 M NaAc and 1% SDS) containing proteinase K (final concentration 0.2 mg/mL, 740506, Macherey-Nagel GmbH & Co KG, Germany). The samples were mixed and incubated overnight at 37°C. Next, RNase A (final concentration 0.2 mg/mL, EN0531, Thermo Fisher Scientific) was added, and the samples were incubated for 30 min at 37°C. Lysis buffer (as described above) and Tris buffered phenol were added to the pellet and after gentle mixing, the samples were centrifuged for 5 min at 4000 rpm at RT. The upper phase was transferred into a new tube, mixed with chloroform-isoamyl alcohol (24:1) and centrifuged for 5 min at 4000 rpm at RT. The pellet was washed twice with 100% ethanol before mtDNA was precipitated from the upper phase by adding 2.5 volumes of 100% ethanol and 0.1 volumes of 3 M NaAc, pH 5.2. The samples were next centrifuged for 30 min at 13 000 x g at 4°C. The pellet was washed with 70% ethanol followed by a 4 min centrifugation at 10 000 x g at RT. The pellet was air dried for 30 min, dissolved into 50 µL of TE buffer (1 M Tris and 0.5 M EDTA, pH 8.0) and heated for 15 min at 37°C in a shaker (500 rpm). RT-qPCR was conducted according to the manufacturer’s protocol with SYBR Green Master Mix (K0252, Thermo Fisher Scientific) and the template amount used was 2 ng per well. The amount of mitochondrial *CYTB*, *DLOOP* and *16S* as well as genomic *HBB*, *APP* and *B2M* were analysed using LightCycler® 480 System (Roche Diagnostics AG, Basel, Switzerland) combined with LightCycler 480 SW 1.5 (Roche Diagnostics AG) (all LM and rosiglitazone related experiments) or the CFX384 Touch Real-Time PCR Detection System combined with CFX Maestro software (Bio-Rad Laboratories Inc., Hercules, CA, USA) (other experiments). The mtDNA amount (mean of *CYTB*, *DLOOP* and *16S* regions) relative to genomic DNA (geometric mean of *HBB*, *APP* and *B2M* regions) was analysed using qBase+ software version 3.4 (Biogazelle, Gent, Belgium). The primers used in the experiments are listed in Table S3.

#### Histology

The adipocyte spheroids were washed twice with 1x DPBS and they were placed on a plastic mold partially filled with pre-frozen Tissue Tek O.C.T. compound (OCT, 45830, Histolab, Products AB, Sweden). This prevented adipocyte spheroids to locate too close to the edges. Before freezing, the adipocyte spheroids were completely embedded in OCT to ensure that adipocyte spheroids were in the center of the mold. Adipocyte spheroids were frozen at -100°C for 3 min using the Minitube Freezer (Minitube GmbH) and stored at -80°C until use. Prior to cutting, the molds were stored at - 25°C overnight to adjust OCT to the cutting temperature. The cryosections with a thickness of 10 μm were cut using the Leica CM3050 cryostat at a temperature of - 25°C to -30°C. The cryosections were then transferred to slides and stored at -20°C until staining. The haematoxylin-eosin (HE, 01820 and 1650, respectively, Histolab, Askim, Sweden) stainings were performed at RT after thawing the cryosections at 4°C for at least 10 min. The samples were fixed for 10 min with acetone at RT. After air drying, the undifferentiated preadipocyte spheroids were stained with haematoxylin for 1 min and all differentiated adipocyte spheroids for 2 min, then cryosections were washed for 5 min with tap water followed by washing with 10 sec in distilled water, before 2 min of eosin staining. Next, the cryosections were washed with increasing ethanol concentrations (3x for 10 sec in 96% ethanol, 1x for 10 sec in 100% ethanol, 1x for 1 min in 100% ethanol and again 2x for 2 min in 100% ethanol) and finally 2x in 100% xylene for 2 min. After adding the embedding medium ROTI®-Histokitt (6638.2, Carl Roth GmbH & Co KG), the cryosections were covered with a coverslip and stored at RT until imaging with PreciPoint M8 digital microscope (PreciPoint GmbH, Freising, Germany) at 20x magnification and viewed with the software ViewPoint Light (PreciPoint GmbH).

#### Statistical analyses

All statistical analyses were conducted using GraphPad Prism software version 10.1.2. First, the outliers were removed using ROUT (Q = 1%) and excluded from the analysis. The normality of distributions was evaluated by the Shapiro–Wilk test. For the two group comparisons, unpaired t-test or nonparametric Mann Whitney U-test was used when appropriate. The three group comparisons were analysed using analysis of variance (ANOVA) for normal distribution or Kruskal–Wallis test for non-normal distribution. ANOVA was followed by the Fisher’s Least Significant Difference (LSD) test, whereas the Kruskal–Wallis test was followed by the Uncorrected Dunn’s test to assess differences of planned comparisons among the groups. The sample sizes of the groups in each specific experiment are indicated in the figure legends. The level of statistical significance was set at *P* < 0.05 and the data are presented as means ± SEM if not stated otherwise.

## RESULTS

### Optimization of the cell culture conditions and differentiation protocol

To create a human adipocyte 3D model for studying mitochondrial function, we started by optimizing differentiation conditions using preadipocytes from two donors with different BMIs (donors 1 and 2, Table 1). The preadipocytes were first expanded in monolayer cultures in growth media supplemented with human serum to avoid exposure to xenobiotic compounds and then transferred to 3D culture plates. Differentiation was induced with commercially available chemically defined differentiation and maturation media devoid of serum. This approach was chosen as human adipose-derived precursor cells and preadipocytes do not require serum for differentiation (14, 15, 20). Moreover, PPARγ agonist thiazolidinedione-free differentiation media was used throughout all experiments to preserve the white adipocyte phenotype upon differentiation.

To find the optimal adipocyte spheroid size and differentiation duration, we seeded 15 000, 25 000 or 35 000 preadipocytes per well and differentiated the spheroids under scaffold-free conditions for 6, 13 or 20 days in differentiation media followed by a 7-day culture in maintenance media (Fig. S1A). This resulted in a 2-, 3- or 4-week-long differentiation protocol, respectively. Subsequent gene expression analysis showed that the largest adipocyte spheroids (35 000 cells) had the most consistent upregulation of adipogenesis marker *PPARγ* after the 3-week differentiation protocol (Fig. S1B). Extending differentiation beyond three weeks did not markedly increase *PPARγ* expression levels in either donor (Fig. S1B). Notably, *PPARγ* expression levels in adipocyte spheroids were very similar to those of subcutaneous MAFs (donor 3, Table 1, Fig. S1B) suggesting that our protocol produces white adipocytes with the physiologically relevant gene expression. Based on these results, we selected 35 000 preadipocytes per spheroid with a 3-week differentiation duration for subsequent experiments.

### Extracellular matrix improves mitochondrial network and respiration

Previous studies have shown that extracellular matrix (ECM) and especially the addition of ECM protein mixtures such as GFR-Matrigel improves lipid droplet morphology and the metabolic phenotype of human white adipocyte spheroids (11, 19). Thus, we determined whether GFR-Matrigel also affects mitochondrial metabolism by using adipocyte spheroids derived from the donor 2. After a 3-week differentiation period, the human adipocyte spheroids were stained with MitoTracker, a mitochondrial marker, and analysed using fluorescence live cell imaging. The results revealed that the scaffold-free adipocyte spheroids had more dense spheroid structure as well as smaller and more fragmented mitochondria (Fig. S2A). In contrast, the GFR-Matrigel embedded adipocyte spheroids exhibited a loose spheroid structure with highly connected and elongated mitochondrial network (Fig. S2A). Bright-field images after a 3-week differentiation period verified the morphological differences; the scaffold-free adipocyte spheroids were smaller and more compact, while the GFR-Matrigel embedded adipocyte spheroids were larger with a visible dense core and a fluffy outer rim, as previously reported for other ECM-embedded adipocyte spheroids (19, 22, 23) (Fig. S2B).

To evaluate how GFR-Matrigel influences mitochondrial function, we established and optimized mitochondrial respiration measurement with the Seahorse Analyzer using adipocyte spheroids derived from the donor 2. Interestingly, differentiation of adipocyte spheroids within GFR-Matrigel significantly improved several mitochondrial respiration-related variables including basal, ATP-linked and maximal respiration as well as the spare respiratory capacity when compared to scaffold-free differentiated adipocyte spheroids (Fig. S2C-D). We did not observe significant changes in proton leak (*i.e.*, uncoupling) or coupling efficiency (*i.e.*, the proportion of mitochondrial electron transport chain (ETC) activity used to drive basal state ATP synthesis) (Fig. S2C-D). We also tested how the cell number per (pre)adipocyte spheroid affects the unnormalized cellular respiration rates *i.e.* OCR values and the reliability of the Seahorse measurements. Notably, unnormalized OCR values above 20 pmol/min have been considered as trustworthy for respiration detection (21). Although basal and maximal respiration OCR values for the GFR-Matrigel embedded differentiated adipocyte spheroids containing 15 000 cells were above 20 pmol/min, the scaffold-free undifferentiated preadipocyte spheroids with the same cell number did not yield reliable OCR values (Fig. S2E). However, the scaffold-free undifferentiated spheroids with 35 000 cells did demonstrate OCR values above 20 pmol/min (Fig. S2E). This suggests that a cell count of 35 000 is required for reliable detection of mitochondrial respiration both in the undifferentiated and differentiated (pre)adipocyte spheroids. Collectively, the embedding of human adipocyte spheroids in GFR-Matrigel seemed to offer an advantageous environment for adipocytes’ mitochondrial network and respiration. As a result, we selected the spheroid formation protocol involving 35 000 preadipocytes embedded in GFR-Matrigel and differentiation in human serum- and PPARγ agonist thiazolidinedione-free media for three weeks as our gold standard protocol and all subsequent experiments in this article were conducted using this protocol if not stated otherwise.

### Differentiated adipocyte spheroids recapitulate the subcutaneous white adipocyte phenotype

After identification of the optimal differentiation protocol for our adipocyte spheroid model, we characterized the morphology, differentiation marker expression and metabolic properties of the undifferentiated and differentiated (pre)adipocyte spheroids derived from the donors 1 and 2 (Fig. 1A). Upon morphological analysis, we noticed that the size of the adipocyte spheroids increased during the first week of differentiation (Fig. 1B-C). Thereafter, the size remained stable for the donor 1, while the adipocyte spheroids derived from the donor 2 showed a slight increase in diameter between week one and two (Fig. 1B-C). On average, the adipocyte spheroid diameter was 2- to 3- fold higher after 3 weeks, nearly 2 mm, regardless of the cell donor (Fig. 1C). Interestingly, total DNA content, a proxy for the cell number, did not significantly differ between the undifferentiated and differentiated (pre)adipocyte spheroids suggesting that cells did not significantly proliferate during the differentiation (Fig. S3A). Despite the rather large size of the differentiated adipocyte spheroids, cryosections did not reveal signs of a necrotic center (Fig. S3B). In the cryosections, the empty spaces, considered as remnants of lipid droplets and GFR-Matrigel, were abundant throughtout the adipocyte spheroids suggesting the presence of differentiated adipocytes also in the core of the adipocyte spheroids (Fig. S3B). In line, the differentiated adipocyte spheroids from either donor did not demonstrate any significant cell death based on the low lactate dehydrogenase (LDH) enzyme activity in the cell culture media (< 20 U/L vs. ≤ 245 U/L expected in human serum based on a previous report (24) and the manufacturer’s instructions, Fig. S3C).

**Figure 1.**
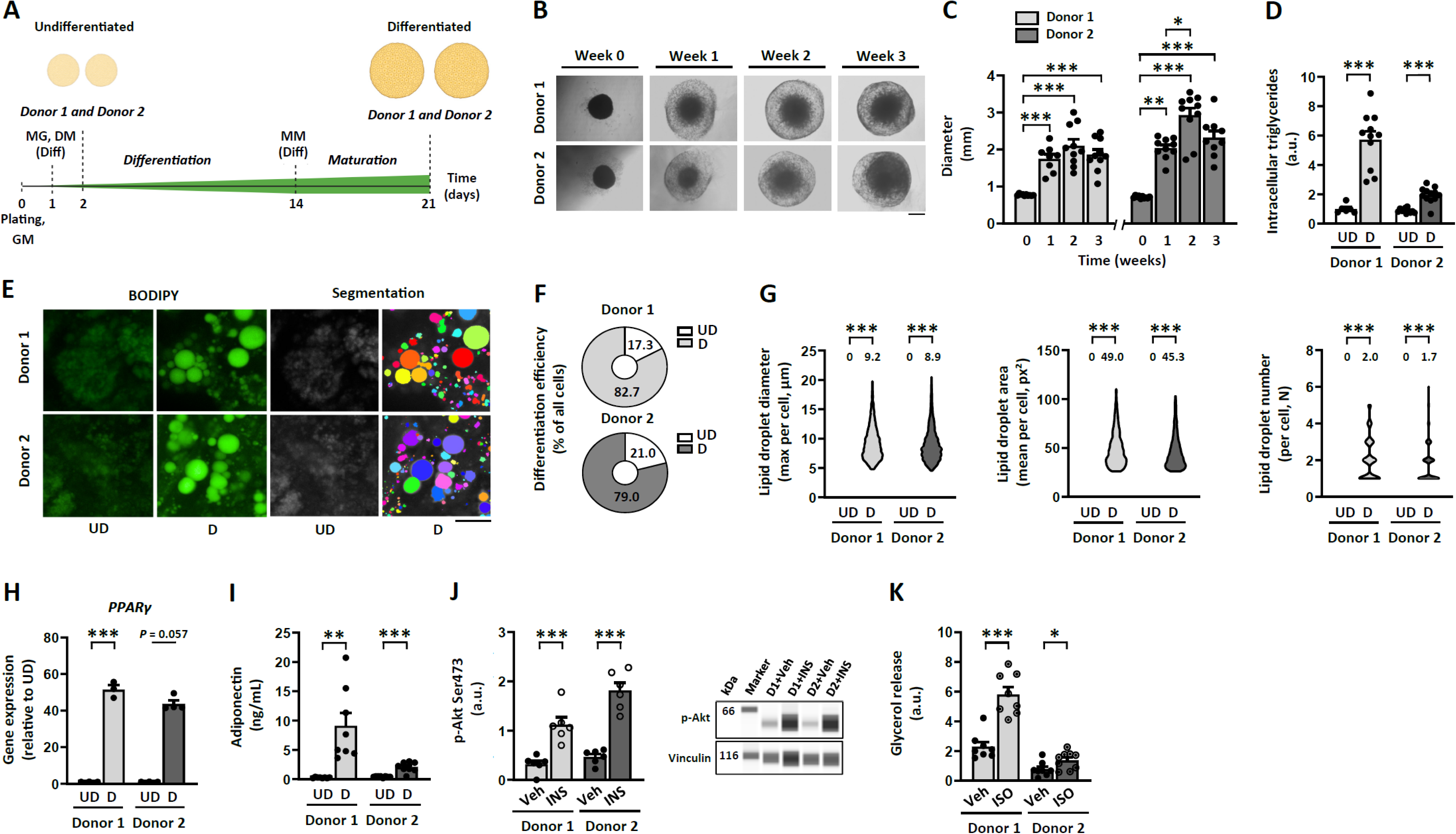
Differentiated human subcutaneous white adipocyte spheroids display a morphological, transcriptional, and metabolic profile typical for mature white adipocytes. A) Schematic presentation of the study design. All experiments were conducted using (pre)adipocyte spheroids derived from the donors 1 and 2. GM, growth media without serum; MG, growth factor reduced-Matrigel; DM, adipocyte differentiation media; Diff, administration to only differentiated adipocyte spheroids; MM, adipocyte maintenance media. B) Selected brightfield images of the adipocyte spheroids from the donors 1 and 2 during the 3-week differentiation period. Scale bar = 500 µm. C) Adipocyte spheroid diameter development during the 3-week differentiation period, N = 8-10 per group. D) Intracellular triglyceride content in undifferentiated (UD) and differentiated (D) (pre)adipocyte spheroids, N = 6-12 per group. E) Selected representative fluorescence live cell images of lipid droplets (BODIPY, green) and the corresponding segmentation masks used for the lipid droplet quantification in differentiated adipocyte spheroids. Scale bar = 20 µm, 40x air objective . F) Differentiation efficiency in differentiated adipocyte spheroids, N = 4 per group. G) Lipid droplet maximal diameter, mean area and number per cell in UD and D (pre)adipocyte spheroids, N = 3-4 (pre)adipocyte spheroids per group. Mean values per group are reported above the violin plots. Note that lipid droplets above the setted threshold were included only from the differentiated adipocyte spheroids. For UD only cells with values below the set threshold were included. The number of cells analyzed are depicted in the brackets for each analysis as follows: lipid droplet maximal diameter (535-2391 cells/spheroid), lipid droplet mean area (537–2625) and lipid droplet number (534-2153). Gene expression of H) *PPAR*γ in UD and D (pre)adipocyte spheroids, N = 3-4 per group. In H), data are presented as relative to UD = 1. I) Media adiponectin concentration from UD and D (pre)adipocyte spheroids, N = 8 per group. J) Protein content of phosphorylated Akt^Ser473^ in response to a 30-min vehicle (Veh, 1x PBS) or 100 nM insulin (INS) administration in differentiated adipocyte spheroids, N = 6 per group. Representative blots are shown on the right. Vinculin was used for data normalization. K) Glycerol release in response to a 3-hour vehicle (Veh, water) or 10 µM isoproterenol (ISO) administration in differentiated adipocyte spheroids, N = 8-9 per group. Data are shown as means with individual values. In C) and J-K), statistical analyses were performed with one-way ANOVA followed by Uncorrected Fisher’s LSD or with Kruskal–Wallis test followed by Uncorrected Dunn’s test. In D) and F-I), unpaired t-test or Mann-Whitney U-test was used. * P < 0.05, ** P < 0.01 and *** P < 0.001.

To validate the level of differentiation and maturation of the adipocytes, we measured several adipogenic markers. The analysis of intracellular triglyceride content showed a significant accumulation of lipids into the differentiated adipocyte spheroids in comparison to the respective undifferentiated preadipocyte spheroids cultured for 2 days (Fig. 1D). The increase in stored triglycerides was higher for the adipocyte spheroids derived from the donor 1 with a lower BMI (Fig. 1D). Next, we used fluorescence live cell imaging with a BODIPY lipid marker to analyze lipid droplet morphology and number in non-fixed human adipocyte spheroids. This technique enabled us to examine large cell population in the outer layer of the adipocyte spheroids. Only in the differentiated adipocyte spheroids, there was an appearance of adipocytes with lipid droplets and these adipocytes demonstrated uni- and multilocular lipid droplet morphology (Fig. 1E, Fig. S4A). The proportion of lipid-containing cells with unilocular lipid droplets was near 50% (donor 1; mean 47.0 ± SEM 5.7% and donor 2; mean 54.3 ± SEM 3.8%) of the analyzed cells per spheroid. The differentiation efficiency, evaluated as the ratio of differentiated adipocytes to the total number of imaged cells per differentiated adipocyte spheroid, was high, nearly 80% regardless of the donor (Fig. 1F). For both donors, the average diameter of the largest lipid droplet per cell (including both unilocular and multilocular droplets) was ∼9 µm for the differentiated adipocyte spheroids (Fig. 1G). Typically, the diameter range varied between 5 to 20 µm (Fig. 1G). On average, the mean lipid droplet area (∼50 px^2^) and lipid droplet number (∼2) per cell were similar in the differentiated adipocyte spheroids from both donors (Fig. 1G). These changes in lipid deposition and lipid droplet morphology were accompanied with upregulated gene expression of common adipogenesis markers, including *PPAR*γ (Fig. 1H), adiponectin (*ADIPOQ*), fatty acid synthase (*FASN*) and hormone sensitive lipase (*HSL*) (Fig. S3D). The changes in gene expression profile were greater in the adipocyte spheroids derived from the donor 1 with a lower BMI (Fig. 1H and Fig. S3D). The gene expression of a beiging/browning marker, *UCP1*, increased in the differentiated adipocyte spheroids in comparison to the respective undifferentiated preadipocyte spheroids, especially in the donor 1 (Fig. S3E). However, in comparison to scWAT and subcutaneous MAF pool from multiple donors (donors 4-9, Table 1), the *UCP1* expression in adipocyte spheroids remained within the normal physiological range (Fig. S3E). In conclusion, our adipocyte spheroids fulfilled the criteria for efficient adipogenic differentiation including elevated lipid accumulation and the formation of white adipocytes with unilocular lipid droplets.

Next, we analyzed adipokine secretion capacity by quantifying media adiponectin concentration. The differentiated adipocyte spheroids from both donors had significantly higher adiponectin secretion as compared to the respective undifferentiated preadipocyte spheroids, but this was more pronounced in the adipocyte spheroids derived from the donor 1 with a lower BMI (Fig. 1I). Interestingly, we noticed that the adiponectin secretion capacity was affected by passaging of predipocytes used to generate the adipocyte spheroids. More precisely, a lower passage number (2-3 vs. 6) of predipocytes markedly facilitated adiponectin secretion to the cell culture media even in the adipocyte spheroids with a slightly lower cell number than in our golden standard protocol (32 500 vs. 35 000 cells per spheroid) (Fig. S3F).

Furthermore, to validate the responsiveness to hormonal signals, we administered the differentiated adipocyte spheroids with insulin or isoproterenol to investigate their downstream protein phosphorylation and lipolytic responses, respectively. Our results showed that independent of the donor charachteristics, insulin stimulation greatly increased the phosphorylation of Akt^Ser473^ in the differentiated adipocyte spheroids (Fig. 1J), while total Akt remained unchanged (Fig. S3G). In addition, the differentiated adipocyte spheroids from both donors responded to isoproterenol by secreting glycerol into the media ≥ 2-fold in comparison to basal, non-stimulated condition (Fig. 1K). That being said, the donor 1 with a lower BMI had a greater response to the isoproterenol administration (Fig. 1K).

Together, these results demonstrated that our adipocyte spheroids exhibit morphological, transcriptional and metabolic profiles typical for *in vivo* subcutaneous white adipocytes (3).

### Differentiated adipocyte spheroids exhibit increased mitochondrial respiration and content

Given that mitochondria are known to be crucial during adipogenesis to fulfill the energy needs of differentiating cells and to support adipokine secretion (25), we next asked whether our adipocyte spheroid model can be exploited to detect the remodelling of mitochondrial metabolism that occurrs during adipocyte differentiation (26, 27). We first used respirometry to compare the undifferentiated and differentiated (pre)adipocyte spheroids from both donors. We found OCR to be in the detectable range with the chosen adipocyte spheroid size (Fig. S3H). The differentiated adipocyte spheroids from both donors showed significantly elevated basal respiration, proton leak, ATP-linked respiration, maximal respiration and spare respiratory capacity in comparison to the respective undifferentiated preadipocyte spheroids (Fig. 2A-B). Coupling efficiency was significantly reduced in the differentiated adipocyte spheroids from both donors in comparison to the undifferentiated preadipocyte spheroids (Fig. 2B), suggesting less efficient mitochondrial respiration after the 3-week differentiation.

**Figure 2.**
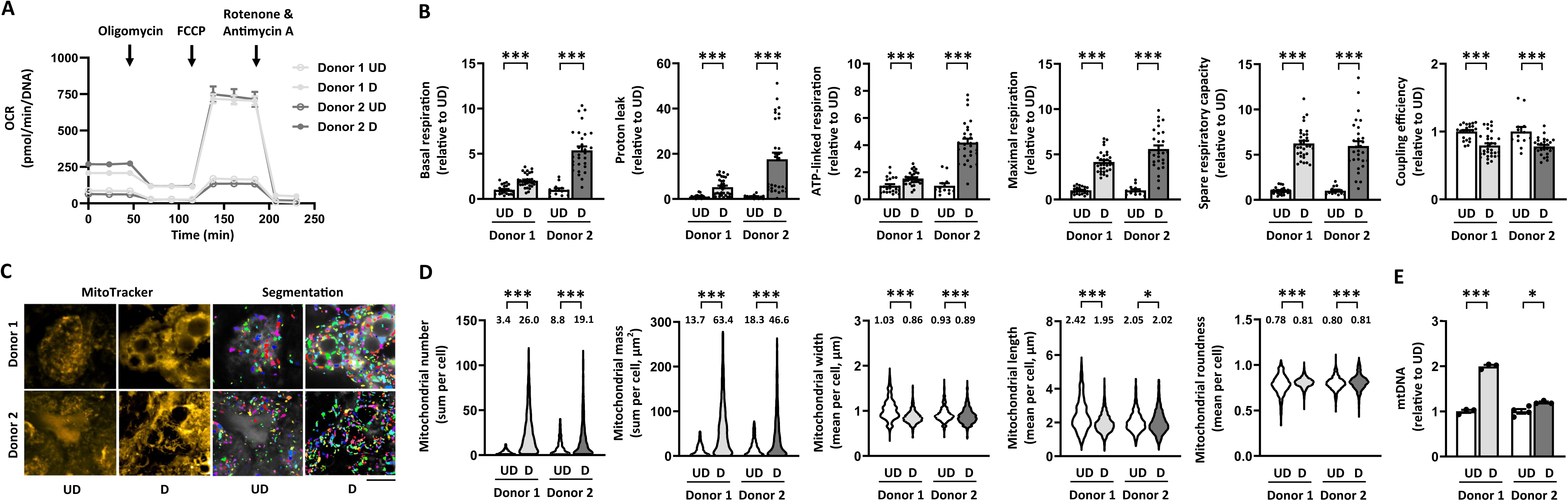
Differentiation of human subcutaneous white adipocyte spheroids increases mitochondrial respiration, biogenesis and fragmentation. A) Mitochondrial respiration analysis in undifferentiated (UD) and differentiated (D) (pre)adipocyte spheroids derived from the donors 1 and 2. Representative oxygen consumption rates (OCR) were normalized to total DNA amount. Parameters of mitochondrial respiration, including B) basal respiration, proton leak, ATP-linked respiration, maximal respiration, spare respiratory capacity and coupling efficiency in UD and D (pre)adipocyte spheroids, N = 13-32 per group. C) Selected representative fluosescence live cell images of the mitochondria (MitoTracker, yellow) and the corresponding segmentation masks used for the quantification of mitochondrial parameters in undifferentiated and differentiated (pre)adipocyte spheroids from the donors 1 and 2. Scale bar = 20 µm, magnification is 40x air objective. Quantification of D) mitochondrial number, mass, width, length and roundness based on the fluorescence live cell imaging, N = 3-4 (pre)adipocyte spheroids per group. Mean values per group are reported above the violin plots. The number of cells analyzed are depicted in the brackets for each analysis as follows: mitochondrial number (648-2960 cells/spheroid), mitochondrial mass (713-2966), mitochondrial width (764-3176), mitochondrial length (747-3184) and mitochondrial roundness (788-3185). E) Mitochondrial DNA (mtDNA) amount expressed per nuclear genome in UD and D (pre)adipocyte spheroids, N = 3-4 per group. In B) and E), data are presented as relative to UD = 1. Data are shown as means with individual values. In B and D), unpaired t-test or Mann-Whitney U-test was used. * P < 0.05 and *** P < 0.001.

To assess changes in mitochondrial content and morphology upon adipocyte spheroid differentiation, we employed fluorescence live cell imaging focusing on the outer layer of adipocytes of the differentiated spheroids (Fig. 2C, Fig. S4A). Upon differentiation, mitochondrial number and mass per cell were significantly increased (Fig. 2D), the donor 1 with a lower BMI demonstrating a greater increase in these two variables. Mitochondrial width and length slightly decreased upon differentiation, while roundness increased, particularly in the donor 1, possibly leading to the formation of smaller, more fragmented mitochondria (Fig. 2D). In line with the imaging data, the amount of mitochondrial DNA (mtDNA) per nuclear genome was significantly increased in the differentiated adipocyte spheroids derived from both donors in comparison to the respective undifferentiated preadipocyte spheroids (Fig. 2E). The increase in mtDNA amount was higher in the adipocyte spheroids derived from the donor 1 with a lower BMI.

Taken together, our results demonstrated that adipocyte spheroid differentiation induced an increase in mitochondrial respiration and biogenesis, as expected based on the growing body of literature (26, 27). These findings highlight the suitability and usability of our human adipocyte spheroid model for reliable analysis of mitochondrial metabolism related variables in physiological conditions.

### Lipid mixture (LM) administration promotes lipid accumulation and impairs metabolic homeostasis and mitochondrial function

To evaluate whether we could mimic mitochondrial dysfunction and metabolic complications in adipocytes typically induced by a nutrient-based energy surplus during obesity (3), we added a lipid mixture (LM) to the media of adipocyte spheroids derived from the donor 2 with a higher BMI during the 3-week differentiation protocol (Table 1, Fig. 3A). We selected a commercial LM that contains both unsaturated and saturated fatty acids as well as cholesterol because everyday foods normally contain a mixture of these lipid classes (28, 29).

**Figure 3.**
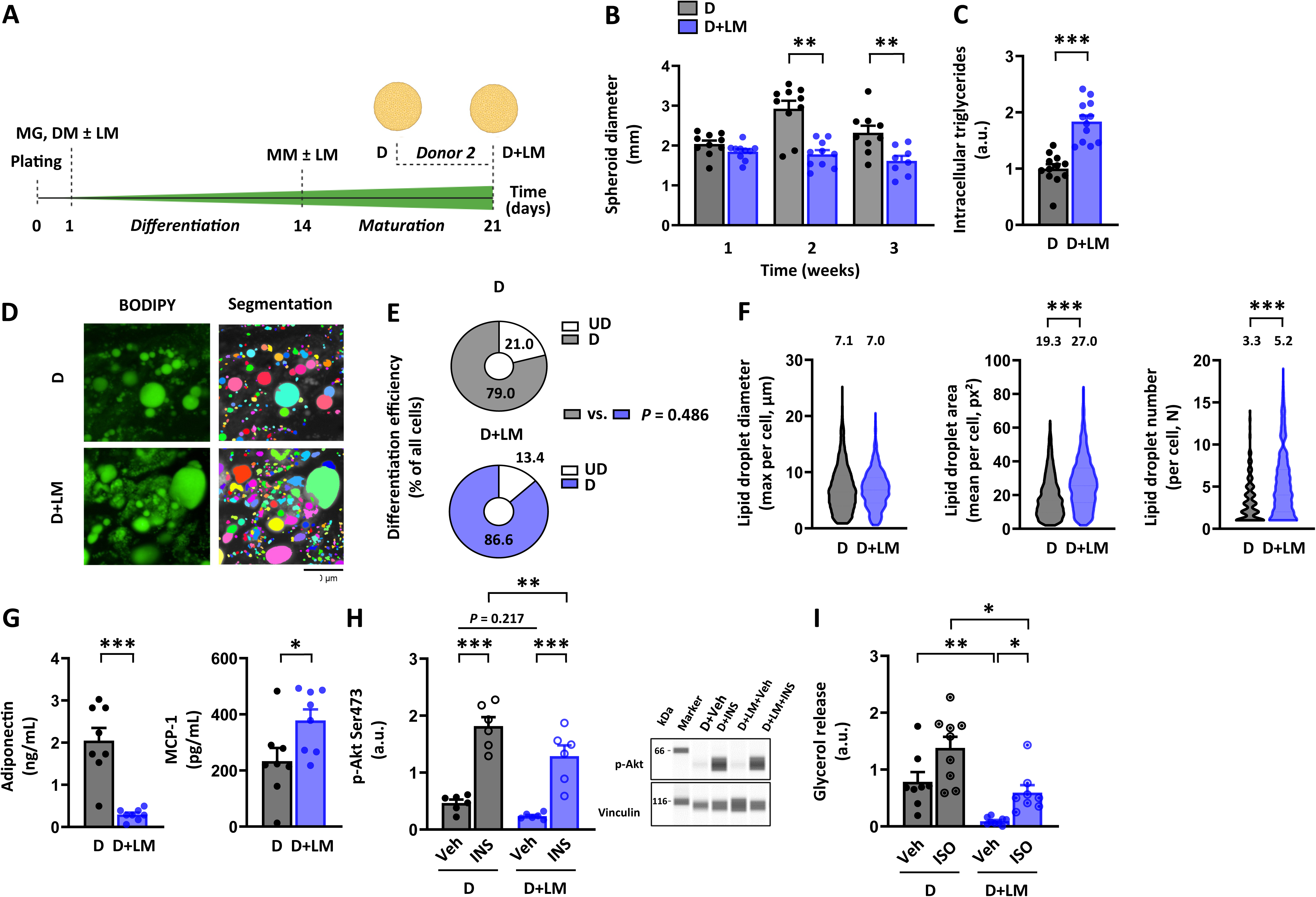
Lipid mixture administration increases lipid accumulation and impairs endocrine function and hormonal responsiveness. A) Schematic presentation of the study design. Only differentiated adipocyte spheroids derived from the donor 2 were used in these experiments. B) Adipocyte spheroid diameter development without (D) or with lipid mixture (LM) administration (D+LM) during the 3-week differentiation period, N = 8-10 per group. C) Intracellular triglyceride content in D and D+LM adipocyte spheroids, N = 12 per group. Data are presented as relative to D = 1. D) Selected representative fluorescence live cell images of lipid droplets (BODIPY, green) and the corresponding segmentation masks used for the lipid droplet quantification in D and D+LM adipocyte spheroids. Scale bar = 20 µm, magnification 40x air objective. E) Differentiation efficiency in D and D+LM adipocyte spheroids, N = 4 per group. F) Lipid droplet maximal diameter, mean area and number in D and D+LM adipocyte spheroids, N = 4 per adipocyte spheroids group. Mean values per group are reported above the violin plots. The number of cells analyzed are depicted in the brackets for each analysis as follows: lipid droplet maximal diameter (2372-4495 cells/spheroid), lipid droplet mean area (3060-4377) and lipid droplet number (1832-4044). Media G) adiponectin and monocyte chemoattractant protein 1 (MCP-1) concentrations from D and D+LM adipocyte spheroids, N = 8 per group. H) Protein content of phosphorylated Akt^Ser473^ in response a 30-min vehicle (Veh, 1x PBS) or 100 nM insulin (INS) administration in D and D+LM adipocyte spheroids, N = 6 per group. Representative blots are shown on the right. Vinculin was used for data normalization. I) Glycerol release in response to a 3-hour vehicle (Veh, water) or 10 µM isoproterenol (ISO) administration in D and D+LM adipocyte spheroids, N = 8-9 per group. Data are shown as means with individual values. In In B-C) and E-G), unpaired t-test or Mann-Whitney U-test was used. In H-I), one-way ANOVA followed by Uncorrected Fisher’s LSD or with Kruskal–Wallis test followed by Uncorrected Dunn’s test was used. * P < 0.05, ** P < 0.01 and *** P < 0.001.

Starting in week 2 of LM administration, the adipocyte spheroid diameter remained ∼50% lower for as compared to the LM-free controls, with a diameter around 1.5 mm per spheroid at the end of the differentiation (Fig. 3B, Fig. S5A). Given that total DNA content decreased for the adipocyte spheroids after LM administration in comparison to the LM-free adipocyte spheroids (Fig. S5B), the smaller adipocyte spheroid size could be due to the lower cell proliferation and/or more efficient packing of lipids into larger lipid droplets. Notably, LM administration did not promote cell death compared to the LM-free counterparts as evaluated based on LDH activity in the media (Fig. S5C).

As expected, intracellular triglyceride content in adipocyte spheroids was significantly increased in response to LM administration during differentiation (Fig. 3C). Based on the fluorescence live cell imaging (Fig. 3D, Fig. S4B) and image analysis, we found that the differentiation efficiency did not significantly differ between the LM-administered and LM-free counterparts in the imaged areas (Fig. 3E). In the LM-administered adipocyte spheroids, the maximum lipid droplet diameter per cell was comparable to that of LM-free counterparts, but both mean lipid droplet area and number per cell were significantly elevated (Fig. 3F). LM administration was associated with a slight decline in the proportion of unilocular cells but it did not reach statistical significance (D; mean 29.0 ± SEM 6.0% and D+LM; mean 19.3 ± SEM 0.8%, *P* = 0.155). Importantly, the expression of *PPAR*γ and *UCP1* did not significantly differ between the LM-administered and LM-free counterparts (Fig. S5D). In conclusion, LM administration increased lipid accumulation and promoted the formation and enlargement of lipid droplets, without affecting adipocyte spheroid differentiation or beiging.

In comparison to the LM-free counterparts, the LM-administered adipocyte spheroids had significantly reduced adiponectin secretion (Fig. 3G). In contrast, the secretion of proinflammatory cytokine MCP-1, which is known to be induced in WAT in obesity (30), was significantly enhanced after LM administration (Fig. 3G). Besides impairing adiponectin secretion, LM-induced lipid accumulation also affected their responsiveness to hormonal stimuli. Namely, insulin-stimulated p-Akt^Ser473^ levels were lower after LM administration, suggesting an impairment of insulin signaling (Fig. 3H). Total Akt remained unchanged in all conditions (Fig. S5E). LM administration also significantly reduced glycerol release in basal and isoproterenol stimulated conditions (Fig. 3I). Thus, LM-administration impaired adipocyte spheroid endocrine secretion and hormonal responsiveness.

The Seahorse mitochondrial respirometry assay revealed that LM administration during differentiation significantly reduced basal respiration, proton leak, ATP-linked respiration, maximal respiration and spare respiratory capacity in the adipocyte spheroids compared to their LM-free counterparts (Fig. 4A-B). Coupling efficiency was also markedly reduced, indicating mitochondrial inefficiency following chronic exposure to LM (Fig. 4B). Fluorescence live cell imaging (Fig. 4C, Fig. S4B) and image analysis showed a slight decrease in mitochondrial number but a significant increase in mitochondrial mass compared to the LM-free counterparts (Fig. 4D). LM administration, while reducing roundness, increased mitochondrial width and length, suggesting possibly larger and more elongated mitochondria (Fig. 4D). Additionally, mtDNA content showed an increasing trend after LM administration (Fig. 4E). These findings suggest that LM administration induces mitochondrial respiration defects alongside increased mitochondrial biogenesis.

**Figure 4.**
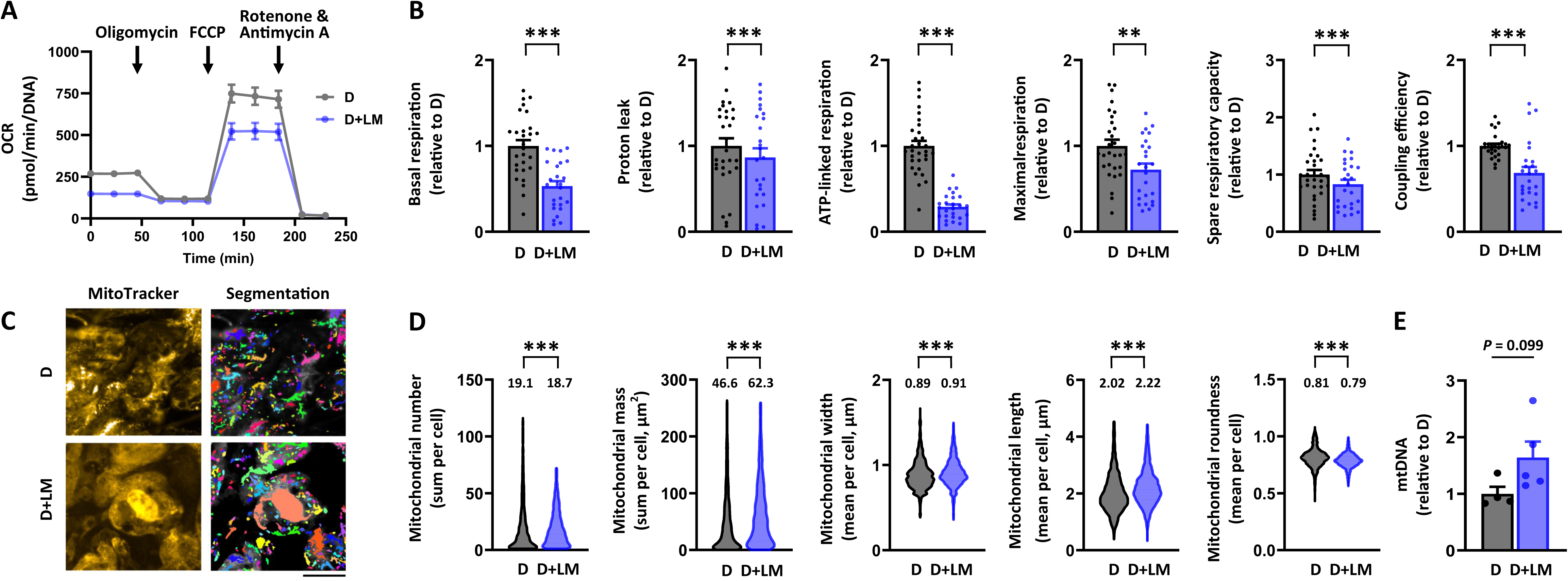
Lipid mixture administration results in mitochondrial bioenergetic defect. Only differentiated adipocyte spheroids derived from the donor 2 were used in these experiments. A) Mitochondrial respiration analyses in differentiated adipocyte spheroids without (D) and with lipid mixture (D+LM) administration. Representative oxygen consumption rates (OCR) were normalized to total DNA amount. Parameters of mitochondrial respiration, including B) basal respiration, proton leak, ATP-linked respiration, maximal respiration, spare respiratory capacity and coupling efficiency in D and D+LM adipocyte spheroids, N = 25-31 per group. C) Selected representative fluorescence live cell imaging of the mitochondria (MitoTracker, yellow) and the corresponding segmentation masks used for the quantification of mitochondrial parameters in D and D+LM adipocyte spheroids. Scale bar = 20 µm, magnification 40x air objective. Quantification of D) mitochondrial number, mass, width, length and roundness based on the fluorescence live cell imaging. Mean values per group are reported above the violin plots. N = 4 adipocyte spheroids per group. The number of cells analyzed are depicted in the brackets for each analysis as follows: mitochondrial number (1759-2473 cells/spheroid), mitochondrial mass (2089-2447), mitochondrial width (2163-2671), mitochondrial length (2157-2659) and mitochondrial roundness (2209-2737). E) Mitochondrial DNA (mtDNA) amount expressed per nuclear genome in D and D+LM adipocyte spheroids, N = 4 per group. In B) and E), data are presented as relative to D = 1. Data are shown as means with individual values. In B and D), unpaired t-test or Mann-Whitney U-test was used. ** P < 0.01 and *** P < 0.001.

Collectively, LM administration during adipocyte spheroid differentiation promoted lipid accumulation, along with lipid droplet biosynthesis and enlargement, and adipocyte dysfunction. These alterations were accompanied by mitochondrial bioenergetic defects and a compensatory increase in mitochondrial biogenesis. We conclude that our adipocyte spheroid model with LM administration can be exploited to mimic and study nutrient-induced obesity-associated mitochondrial dysfunction and metabolic disturbances.

#### Rosiglitazone alleviates LM administration-induced disturbances in mitochondrial metabolism

After demonstrating that our adipocyte spheroids serve as a viable model for studying adipocyte mitochondrial metabolism also under pathophysiological conditions, we next aimed to determine whether the adverse effects of LM administration on mitochondrial function and metabolic homeostasis and could be mitigated or even reversed pharmacologically. For this purpose, we chose to use the antidiabetic drug rosiglitazone, which activates PPARγ-controlled transcription (31). In these experiments, LM-administered adipocyte spheroids derived from the donor 2 were treated with rosiglitazone at the concentration of 1 μM for 72 hours after first undergoing the 3-week differentiation protocol (+LM+R, Fig. 5A). As PPARγ activation is known to upregulate transcription of its downstream target genes, such as fatty acid translocase (*FAT*, also known as *CD36*), liver X receptor α (*LXRα*) and adiponectin (*ADIPOQ*) (31, 32), we used expression of these genes as a read-out for rosiglitazone-induced PPARγ activation. We observed that LM administration combined with vehicle treatment (+LM+Veh) significantly downregulated the expression of *CD36*, *LXRα* and *ADIPOQ* in comparison to vehicle without LM (-LM+Veh), which was rescued by rosiglitazone treatment (Fig. 5B). *PPARγ* expression was significantly downregulated by LM administration both in combination with vehicle and with rosiglitazone (Fig. S5F). Notably, *UCP1* expression remained unchanged after rosiglitazone treatment together with LM (Fig. S5F). Collectively, these results confirmed that rosiglitazone treatment activated PPARγ transcriptional activity in our differentiated adipocyte spheroids without affecting adipocyte beiging.

**Figure 5.**
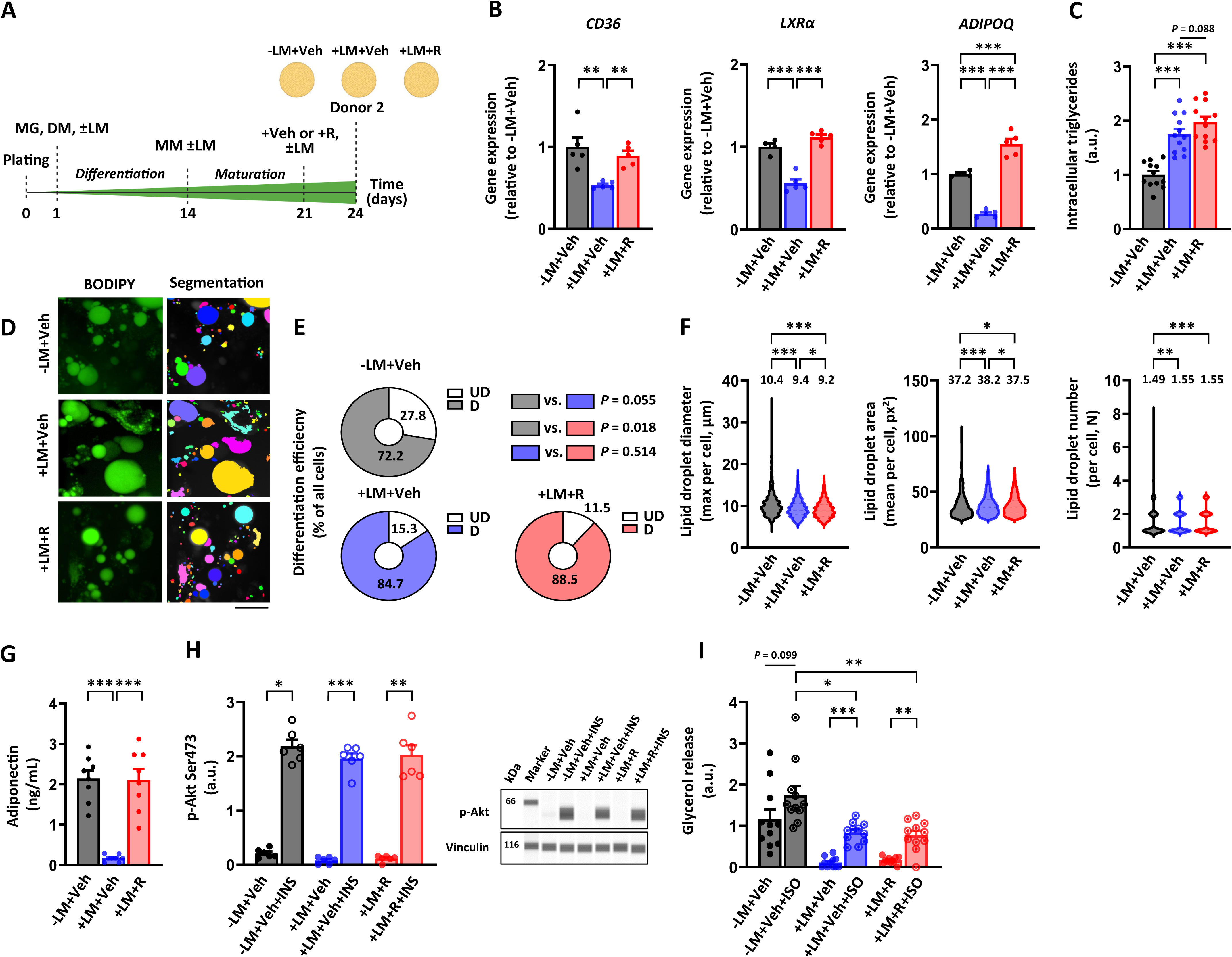
Rosiglitazone treatment increases PPARγ transcriptional activity and restores adipokine secretion with a lesser effect on hormonal responsiveness. A) Schematic presentation of the study design. Only differentiated adipocyte spheroids derived from the donor 2 were used in these experiments. Gene expression of B) *CD36*, *LXRα* and *ADIPOQ* in differentiated adipocyte spheroids administered without lipid mixture (LM) but with vehicle (-LM+Veh), with LM and vehicle (+LM+Veh) or with LM and 1 µM rosiglitazone (+LM+R) for 72 hours post-differentiation, N = 4-5 per group. Data are presented as relative to -LM+Veh = 1. C) Intracellular triglyceride content in adipocyte spheroids between the groups, N = 12 per group. D) Selected representative fluorescence live cell images of lipid droplets (BODIPY, green) and the corresponding segmentation masks used for the lipid droplet quantification in adipocyte spheroids from the different groups. Scale bar = 20 µm, magnification 40x air objective. E) Differentiation efficiency in adipocyte spheroids among the groups, N = 4 per group. The results of statistical analyses between the groups are presented on the top right. F) Lipid droplet maximal diameter, mean area and number in adipocyte spheroids between the groups, N = 4 adipocyte spheroids per group. Mean values per group are reported above the violin plots. The number of cells analyzed per group are depicted in the brackets for each analysis as follows: lipid droplet maximal diameter (882-2234 cells/spheroid), lipid droplet mean area (835-2085) and lipid droplet number (832-2064). G) Adiponectin concentration between the groups, N = 8 per group. H) Protein content of phosphorylated Akt^Ser473^ in response to a 30-min vehicle (Veh, 1x PBS) or 100 nM insulin (INS) administration in adipocyte spheroids between the groups, N = 6 per group. Representative blots are shown on the right. Vinculin was used for data normalization. I) Glycerol release in response to a 3-hour vehicle (Veh, water) or 10 µM isoproterenol (ISO) administration in adipocyte spheroids between the groups, N = 11 per group. Data are shown as means with individual values. In B-C) and E-I), one-way ANOVA followed by Uncorrected Fisher’s LSD or with Kruskal– Wallis test followed by Uncorrected Dunn’s test was used. * P < 0.05, ** P < 0.01 and *** P < 0.001.

Rosiglitazone treatment did not significantly promote intracellular triglyceride accumulation into LM administered adipocyte spheroids from the donor 2 as compared to vehicle (Fig. 5C). Based on the fluorescence live cell imaging (Fig. 5D, Fig. S4C) and image analysis, we found differentiation efficiency to be enhanced in both vehicle and rosiglitazone-treated LM-administered adipocyte spheroids compared to their LM-free counterparts (Fig. 5E). However, rosiglitazone did not significantly impact the differentiation efficiency within the LM-administered adipocyte spheroids (Fig. 5E). Rosiglitazone exacerbated the LM-induced decline in the maximal diameter of lipid droplets but it mitigated the LM-induced increase in the mean lipid droplet area (Fig. 5F). In turn, rosiglitazone did not influence the increase in lipid droplet number caused by LM administration (Fig. 5F). The fraction of unilocular cells in adipocyte spheroids was not significantly affected by rosiglitazone treatment as compared to vehicle (-LM+Veh; mean 69.0 ± SEM 10.6%, +LM+Veh; mean 57.3 ± SEM 2.8% and +LM+R; mean 56.8 ± SEM 4.0%, +LM+Veh vs. +LM+R, *P* = 0.959). These results suggest that rosiglitazone may promote the shrinkage of the lipid droplet size in adipocyte spheroids.

Interestingly, rosiglitazone treatment completely restored the LM-induced decrease in adiponectin secretion from the adipocyte spheroids derived from the donor 2 (Fig. 5G). In contrast, basal and insulin-stimulated Akt^Ser473^ phosphorylation were not markedly affected by rosiglitazone treatment upon LM administration (Fig. 5H). In addition, total Akt remained unchanged in all analyzed conditions (Fig. S5G). In line with the insulin signaling-related results, rosiglitazone did not alleviate the LM-induced impairment in basal and isoproterenol-stimulated glycerol release in adipocyte spheroids (Fig. 5I).

Notably, rosiglitazone treatment did not improve basal respiration, although it partially corrected the LM-induced defects in ATP-linked respiration, maximal respiration and spare respiratory capacity in the adipocyte spheroids derived from the donor 2 (Fig. 6A-B). In addition, rosiglitazone tended to decrease the LM-induced proton leak with the concomitant increase in coupling efficiency, thus suggesting improved ATP production (Fig. 6B). Fluorescence live cell imaging (Fig. 6C, Fig. S4C) and subsequent image analysis showed that rosiglitazone treatment further amplified the LM-induced increase in mitochondrial number and mass (Fig. 6D). Additionally, while rosiglitazone did not alter mitochondrial length, it further reduced the LM-induced decrease in width, resulting in slightly less round and probably a bit more elongated mitochondria (Fig. 6D). However, rosiglitazone did not affect mtDNA amount (Fig. 6E). Collectively, rosiglitazone likely improved mitochondrial bioenergetics, biogenesis and dynamics post-differentiation in adipocyte spheroids.

**Figure 6.**
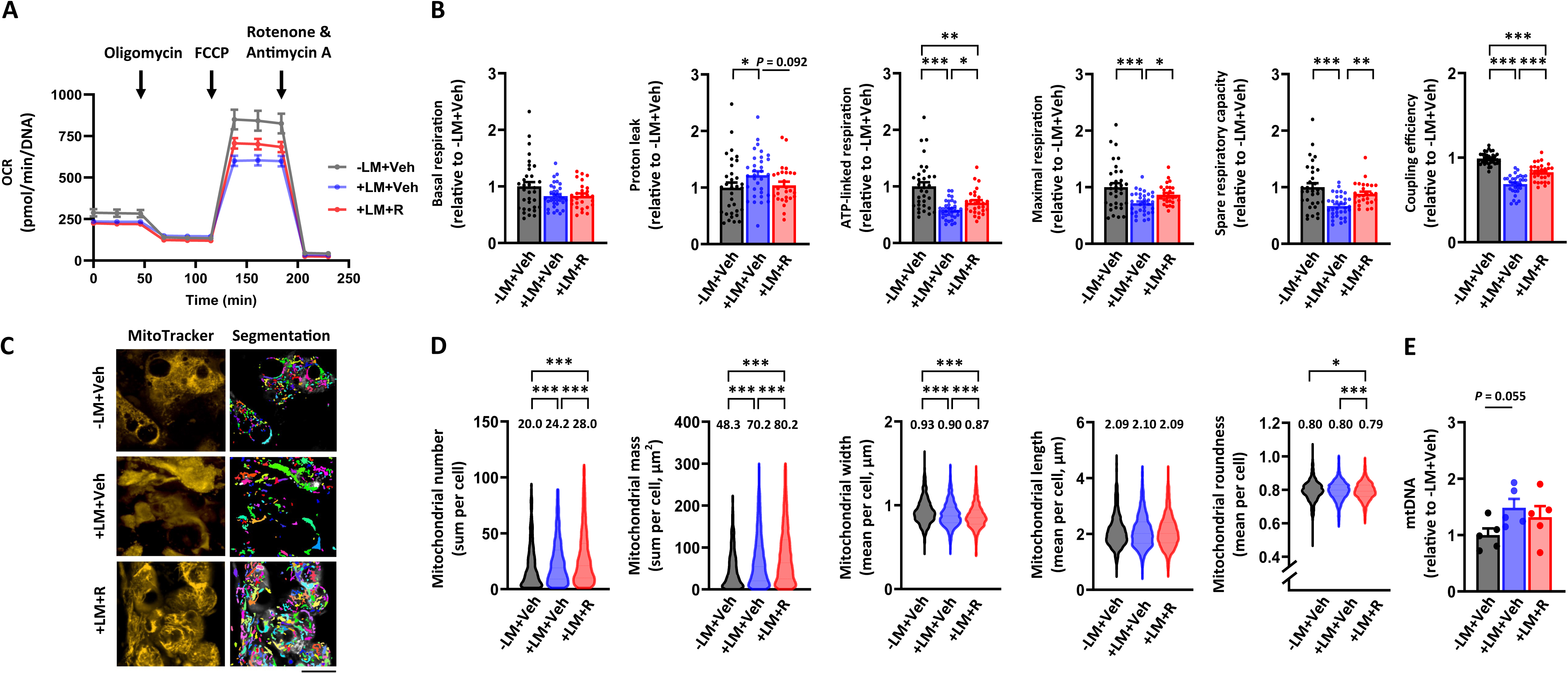
Rosiglitazone treatment alleviates lipid mixture administration-induced disturbances in mitochondrial metabolism. Only differentiated adipocyte spheroids derived from the donor 2 were used in these experiments. A) Mitochondrial respiration analyses in differentiated adipocyte spheroids administered without lipid mixture (LM) but with vehicle (-LM+Veh), with LM and vehicle (+LM+Veh) or with LM and 1 µM rosiglitazone (+LM+R) for 72h post differentiation. Representative oxygen consumption rates (OCR) were normalized to total DNA amount. Parameters of mitochondrial respiration, including B) basal respiration, proton leak, ATP-linked respiration, maximal respiration, spare respiratory capacity and coupling efficiency in adipocyte spheroids between the groups, N = 28-33 per group. Selected representative fluorescence live cell images of mitochondria (MitoTracker, yellow) and the corresponding segmentation masks used for the quantification of mitochondrial parameters in adipocyte spheroids between the groups. Scale bar = 20 µm, magnification 40x air objective. Quantification of D) mitochondrial number, mass, width, length and roundness were analysed based on the imaging analysis. Note that for clarity, y-axis in mitochondrial roundness does not start from zero. Mean values per group are reported above the violin plots, N = 4 adipocyte spheroids per group. The number of cells analyzed per group are depicted in the brackets for each analysis as follows: mitochondrial number (1653-3293 cells/spheroid), mitochondrial mass (1654-3370), mitochondrial width (1790-3414), mitochondrial length (1748-3415) and mitochondrial roundness (1807-3476). E) Mitochondrial DNA (mtDNA) amount expressed per nuclear genome in adipocyte spheroids, N = 5 per group. In B) and E), data are presented as relative to -LM+Veh = 1. Data are shown as means with individual values. In B and D), one-way ANOVA followed by Uncorrected Fisher’s LSD or with Kruskal–Wallis followed by Uncorrected Dunn’s test was used. * P < 0.05, ** P < 0.01 and *** P < 0.001.

As a whole, our results demonstrated that rosiglitazone treatment increased PPARγ transcriptional activity, which was followed by first-line improvements in mitochondrial metabolism, adiponectin secretion, and lipid droplet size. Our findings underscore that our adipocyte spheroid model has the potential to be used in pharmacological studies aimed at improving mitochondrial activity, metabolic homeostasis and lipid droplet remodelling in human subcutaneous adipocytes.

## DISCUSSION

Accumulating evidence suggests that human adipocyte spheroid models offer more physiologically relevant *in vitro* cultures than traditional 2D monolayer cultures for adipocyte studies (9, 11, 12). However, current adipocyte spheroid models have not been optimized for studying mitochondrial metabolism. Given the crucial role of WAT mitochondrial dysfunction in the development of obesity-related metabolic complications, there is a clear need to explore adipocyte mitochondrial biology and pharmacological modulators in a suitable model. In this study, we present a novel 3D *in vitro* model of human subcutaneous adipocytes to investigate mitochondrial related variables. Our findings demonstrate that this new model facilitates the study of mitochondria during adipogenic differentiation, upon lipid overload-induced mitochondrial dysfunction, and the effects of mitochondria-targeting compounds.

At present, several human 3D adipocyte models exist based on different cell culture techniques and protocols, cultured with or without ECM scaffolds (9, 11, 12). Our model builds on this existing knowledge, sharing features such as the use of human subcutaneous preadipocytes, ultra-low-attachment plates, and a GFR-Matrigel scaffold. The key distinctions of our model include a shorter 3-week differentiation period and larger adipocyte spheroid size (35 000 cells), the latter of which allows robust detection of mitochondria-related parameters. By using our newly established model, we observed adipogenesis-induced mitochondrial metabolic remodeling, including increased mitochondrial biogenesis and a metabolic shift towards a more oxidative energy profile, as seen in monolayer cultures (25–27). Despite this, the mitochondria became more fragmented and less efficient by the end of differentiation. At first glance, these findings may seem surprising. However, they align with previous mouse adipocyte studies, which show that mitochondrial respiration is initially tightly coupled to ATP synthesis during adipogenesis but becomes uncoupled as adipocytes mature, focusing more on fat storage (33). Overall, our model is the first for which a comprehensive analysis of mitochondria-related variables under (patho)physiological conditions has been demonstrated, advancing human adipocyte spheroid research.

The existing 3D adipocyte models address many limitations of monolayer cultures, showing improved differentiation and better metabolic functionality (11, 12). In our adipocyte spheroid model, we achieved a high differentiation efficiency of nearly 80%, compared to 20-50% in monolayer cultures (13). Succesfull differentiation was confirmed by increased lipid deposition and high expression of adipogenic genes (*PPARγ*, *ADIPOQ*, *FASN* and *HSL*), consistent with other 3D models (9, 11, 12). Our adipocyte spheroids exhibited typical features of mature white adipocytes, including low *UCP1* expression, rigid responses to insulin and β-adrenergic stimuli, and high adiponectin secretion. Notably, higher passage numbers (6) reduced adiponectin secretion nearly ten-fold compared to lower passages (2–3). This phenomenon may be related to lower differentiation efficiency or less well differentiated cells upon passaging. Importantly, the adipocyte spheroids with low passage number secreted adiponectin to the media in similar quantities as observed in other 3D culture models (9, 11). In line with the previous knowledge on human adipose stem cell function (34), we identified that other potential factors influencing differentiation are dependent on donor BMI and/or genetic background as phenotypic changes associated with adipogenesis were less evident in the adipocyte spheroids derived from the donor 2 with higher BMI and a different genome than the donor 1.

The 3D adipocyte cultures also display a higher proportion of unilocular lipid droplets than in monolayer cultures. In our model, quantification revealed that 30-50% of adipocytes in the outer layer of the spheroids contained unilocular lipid droplets. Not surprisingly, based on the fluorescence live cell imaging, the average maximal lipid droplet diameter (9 µm) in our adipocyte spheroids was smaller than *in vivo* subcutaneous white adipocytes (20 to 150 µm) (35) but comparable to other similar 3D adipocyte models (19, 20), where the typical droplet diameter has been around 10 µm. Nonetheless, differences in imaging techniques and analysis methods make direct comparison with previously published adipocyte spheroid cultures challenging. For example, the present study used live imaging to quantify all imaged cells (thousands), whereas previous studies (19, 20) employed fixed cell imaging with a maximum of a few hundred cells. So far, the largest lipid droplets (20 to 40 µm) have been seen in vascularized adipocyte spheroids (19), suggesting that the absence of endothelial cells in our model may explain the slightly smaller lipid droplet size. As the cell composition in 3D cultures seems to greatly influence adipocyte lipid droplet maturation, this calls for a need of the development of *in vitro* culture models in which all major cellular components of scWAT are present. Overall, a significant portion of the adipocytes in our adipocyte spheroids exhibit unilocular lipid droplets, with the size comparable to that observed in most other 3D models of adipocytes.

Administration of fatty acids or triglycerides has been shown to induce lipid deposition and metabolic complications in human adipocytes in 3D culture models (19, 20, 36). Consistent with these findings, the addition of LM (a mixture of unsaturated and saturated fatty acids as well as cholesterol) increased the lipid droplet mean size in our adipocyte spheroids and likely stimulated the formation of new lipid droplets, as evidenced by a higher lipid droplet number per cell. While the proportion of unilocular lipid droplets did not significantly change, the LM-induced lipid droplet formation likely occurred in multilocular adipocytes. These changes in lipid droplet morphology were linked to early inflammatory signs, such as elevated levels of the pro-inflammatory cytokine MCP-1, as well as impaired insulin signaling and lipolysis, and endocrine failure. Notably, LM administration during adipogenic differentiation led to significant declines in mitochondrial respiration variables, despite increased mitochondrial mass and fusion. This suggests that LM administration may trigger mitochondrial biogenesis and fusion in response to functional decline, similar to compensatory mechanisms observed in mitochondrial disorders (37). Given that decreased mitochondrial respiration, biogenesis, and fusion are also seen in obesity *in vivo* (3), our *in vitro* results indicate that mitochondrial respiration failure may be an early defect associated with lipid overload. The relationship between elevated MCP-1 levels and mitochondrial dysfunction warrants further investigation to determine causality, as cells with defective mitochondria can produce pro-inflammatory cytokines (38). Overall, these results highlight the relevance of our human adipocyte spheroids as a model system that mimics human subcutaneous white adipocytes in obesity-related conditions and provides a platform for studying the molecular mechanisms of adipocytes’ mitochondrial dysfunction.

As scWAT mitochondrial dysfunction is involved in the pathogenesis of obesity (3), pharmacological compounds that improve adipocytes’ mitochondrial function represent important treatment prospect for obesity. Rosiglitazone has been shown to enhance mitochondrial respiration, biogenesis, and fusion in human white adipocytes (39, 40), but its clinical use is limited due to side effects such as heart failure, weight gain, and peripheral oedema (41). In our adipocyte spheroid model, rosiglitazone increased PPARγ activity, likely leading to beneficial effects on mitochondrial respiration, coupling efficiency, number, mass, and fusion. We also observed a significant reduction in lipid droplet size, which may result from enhanced mitochondrial oxidative metabolism, not beiging, as we did not detect *UCP1* upregulation after treatment. This aligns with previous findings indicating that mitochondrial oxidation of fatty acids derived from lipid droplets requires mitochondrial fusion (42), suggesting a close relationship between lipid droplet metabolism and mitochondrial dynamics. Improvements in mitochondrial outcomes were accompanied with a restored capacity for adiponectin secretion. This is consistent with existing literature linking mitochondrial function to adiponectin secretion (43), and the *ADIPOQ* gene being as a known downstream target of PPARγ (31). While rosiglitazone is known for its insulin-sensitizing effects, no such effect was seen in our Akt phosphorylation analysis. This indicates that the insulin-sensitizing properties of rosiglitazone may operate independently of enhanced insulin signaling via Akt, as previously suggested (44), warranting further investigation of its insulin-sensitizing effects in our adipocyte spheroid model. Overall, our study demonstrates that the first positive outcomes of rosiglitazone treatment include improved mitochondrial metabolism, reduced lipid droplet size and restored adiponectin secretion. These findings also underscore our adipocyte spheroid model as a valuable tool for studying mitochondrial boosters and, although not investigated in the present study, assessing potential mitochondrial toxicity in drug development.

In conclusion, we have developed a new *in vitro* model of human adipocyte spheroids which will allow us and others to investigate human white adipocyte mitochondrial biology in physiological and pathophysiological conditions. Furthermore, our model can potentially serve as an *in vitro* drug testing platform in the anti-obesity drug discovery field. A better understanding of the molecular mechanisms behind adipocyte mitochondrial dysfunction, along with the identification of mitochondria activating compounds, could provide tools for mitochondrial pharmacotherapies and the treatment of obesity-related metabolic disorders.

## LIMITATIONS OF THE STUDY

In this study, we mainly used commercially available preadipocytes to increase model and data transparency. However, despite this benefit, the use of these preadipocytes from two middle-aged healthy female donors, one of whom was normal weight and the other overweight, was also a limitation. We recognize that more studies are needed in the future to clarify the influence of donor characteristics *e.g.*, BMI, sex, age and metabolic diseases, such as diabetes, on the differentiation and functionality of adipocyte spheroids. Moreover, as our adipocyte spheroids were generated using only preadipocytes, the absence of other cell types typically present in scWAT likely influenced our results. For example, due to the absence of other cell types, the lipid droplet morphology and metabolic profile of our adipocyte spheroids may not fully recapitulate those of *in vivo* mature white adipocytes.

Another limitation of our model is the use of GFR-Matrigel, the only animal-derived component in our model, which may induce some non-physiological signals. However, as compared to other scaffolds, only GFR-Matrigel has been proven to provide an optimal ECM stiffness and composition for adipogenic differentiation in human 3D adipocyte cultures (11). One of the disadvantages of GFR-Matrigel is uneven and poorly controlled incorporation into adipocyte spheroids, which increases experimental variation. To overcome this GFR-Matrigel related issues, a high number of replicates are recommended for each experiment.

Lastly, current fluorescence live cell imaging techniques do not allow the imaging of the whole adipocyte spheroids throughout their entire height and length. Consequently, we have been able to acquire information related to the differentiation efficiency and lipid droplet and mitochondrial morphology only from the outer layer of the adipocyte spheroids. However, cryosectioning revealed the presence of differentiated adipocytes also in the core of the adipocyte spheroids. In any case, we present live images of human adipocyte spheroids which are likely to provide more biologically relevant data than images from fixed adipocytes.

## AUTHOR CONTRIBUTIONS

AW, AI, CEH and EP conceptualized the model, AW, JHL-K, KP, AH, MK, LR, AI, SM, SS, SL, TR, SK AO and EP performed experiments and/or analyzed data, AW, JHL-K, KP and EP wrote the manuscript, AW, JHL-K, AH and IR prepared the figures and tables, KH, EH, PE, OU, MM, HP, KHP, MK, KAV, CEH and EP provided resources, all co-authors critically reviewed, and edited the manuscript. All authors approved the final version of the manuscript.

## ACKNOWLEDGEMENTS

We would like to express our gratitude to Tilkka Hospital, Helsinki, Finland for their assistance in tissue sample collection and preparation. Imaging was performed at the Biomedicum Imaging Unit, and FIMM High Content Imaging and Analysis unit services of the University of Helsinki, supported by the Helsinki Institute of Life Science (HiLIFE) and Biocenter Finland. These core facilities were supporting imaging and data analysis. We also thank the Chair of Livestock Biotechnology Unit at TUM for their help in freezing spheroids, and the Chair of Nutrition and Immunology at TUM for their assistance with spheroid processing for imaging and cryosectioning. We extend our thanks to Nick Howe from Agilent for his invaluable help in establishing the Seahorse protocol for adipocyte spheroids. Additionally, we acknowledge Minna Eriksson and Emma Paasikivi for their technical support, Riikka Jokinen for providing materials and Juha Hulmi for analytical support. During the preparation of this article the authors used artificial intelligence to improve readability and check grammar.

