## Supplemental Tables 1&3 and Figures 1-5 for "*In vitro* model of human subcutaneous adipocyte spheroids for studying mitochondrial dysfunction and mitochondria activating compounds"

**Supplementary material for the article entitled "*In vitro* model of human subcutaneous adipocyte spheroids for studying mitochondrial dysfunction and mitochondria activating compounds" by Wagner et al. 2025.**

**Supplementary Table 1**. The primers used for gene expression analyses.

| **Target gene** | **Forward sequence (5´→ 3`)** | **Reverse sequence (5´→ 3`)** |
|---|---|---|
| *ADIPOQ* | TGGTGAGAAGGGTGAGAA | AGATCTTGGTAAAGCGAATG |
| *CD36* | CAGGTCAACCTATTGGTCAAGCC | GCCTTCTCATCACCAATGGTCC |
| *FASN* | CCGAGACACTCGTGGGCTA | CTTCAGCAGGACATTGATGCC |
| *HSL* | AGCCTTCTGGAACATCACCGAG | TCGGCAGTCAGTGGCATCTCAA |
| *IPO8* | AAGAAACCGCGCTTGAGGGG | ATCCTCGCTGAGTGGTGCCA |
| *LXRα* | TGGACACCTACATGCGTCGCAA | CAAGGATGTGGCATGAGCCTGT |
| *PPARγ* | TACTGTCGGTTTCAGAAATGCC | GTCAGCGGACTCTGGATTCAG |
| *UCP1* | CAAATCAGCTCCGCCTCTCT | AATGAATACTGCCACTCCTCCAG |

**Supplementary Table 2 (Excel file)**. Pipeline for fluorescence live cell imaging.

**Supplementary Table 3**. The primers used to amplify mitochondrial DNA (mtDNA) or genomic DNA (gDNA) regions.

| **Target gene** | **Forward sequence 5´→3´** | **Reverse sequence 5´→3´** |
|---|---|---|
| *16S* (mtDNA) | GGGGCGACCTCGGAGCAGAA | ATAGCGGCTGCACCATCGGGA |
| *CYTB* (mtDNA) | GCCTGCCTGATCCTCCAAAT | AAGGTAGCGGATGATTCAGCC |
| *DLOOP* (mtDNA) | CATCTGGTTCCTACTTCAGGG | CCGTGAGTGGTTAATAGGGTG |
| *APP* (gDNA) | TGTGTGCTCTCCCAGGTCTA | CAGTTCTGGATGGTCACTGG |
| *B2M* (gDNA) | TGCTGTCTCCATGTTTGATGTATCT | TCTCTGCTCCCCACCTCTAAGT |
| *HBB* (gDNA) | CAGGTACGGCTGTCATCAGTTAG | CATGGTGTCTGTTTGAGGTTGCT |

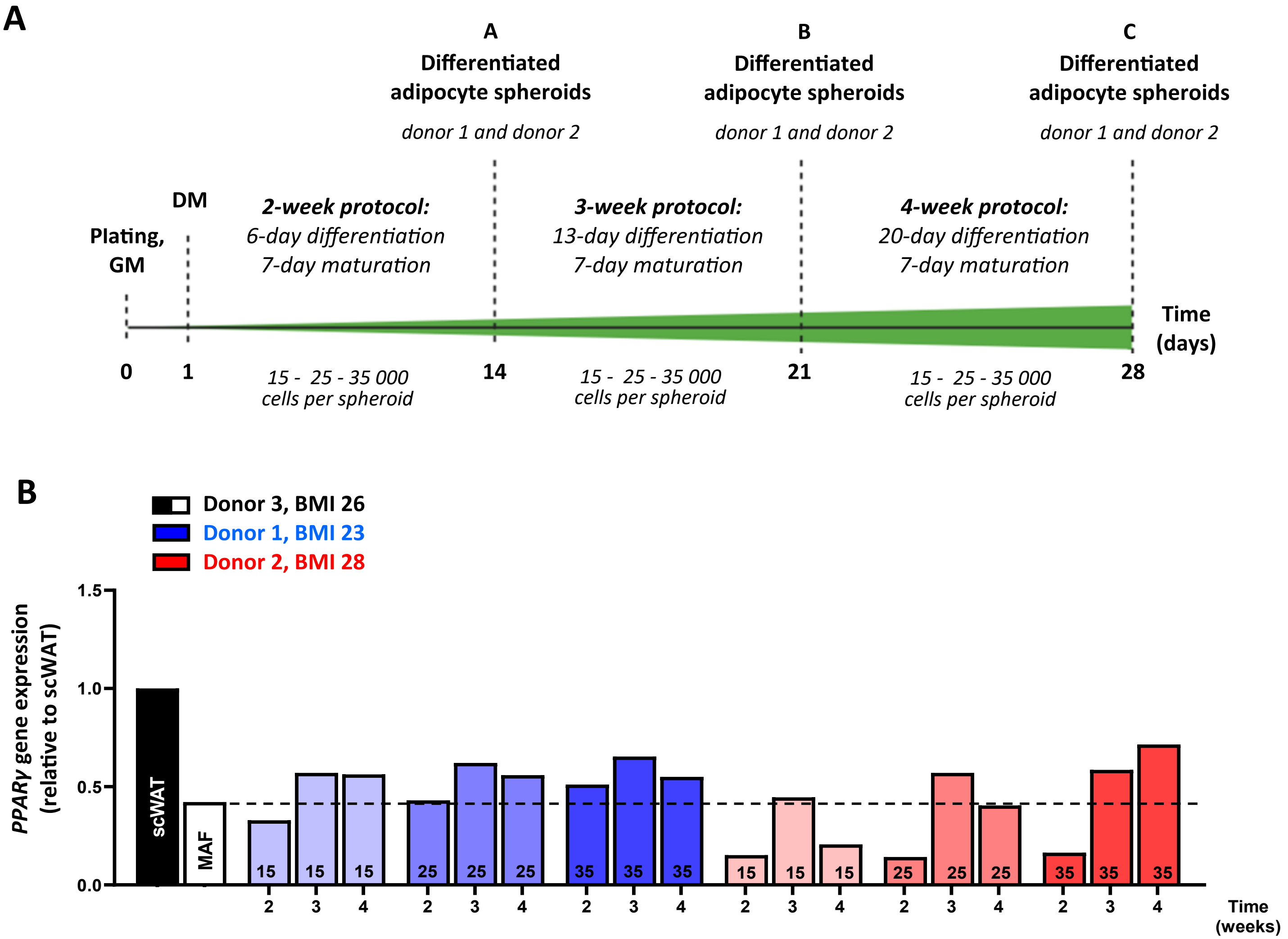
A
A
Differentiated adipocyte spheroids
donor 1 and donor 2
B
Differentiated adipocyte spheroids
donor 1 and donor 2
C
Differentiated adipocyte spheroids
donor 1 and donor 2
Plating, GM
DM
2-week protocol:
6-day differentiation
7-day maturation
3-week protocol:
13-day differentiation
7-day maturation
4-week protocol:
20-day differentiation
7-day maturation
Time (days)
0
1
14
21
28
15 - 25 - 35 000 cells per spheroid
15 - 25 - 35 000 cells per spheroid
15 - 25 - 35 000 cells per spheroid
B
Donor 3, BMI 26
Donor 1, BMI 23
Donor 2, BMI 28
PPARγ gene expression (relative to scWAT)
1.5
1.0
0.5
0.0
scWAT
MAF
15
25
35
2
3
4
Time (weeks)

**Supplementary Figure 1. Validation of the differentiation time and cell number for human subcutaneous white adipocyte spheroid model.** A) Schematic presentation of the study design. Gene expression of B) *PPARγ* in differentiated adipocyte spheroids of various sizes (15 000, 25 000 or 35 000 cells per spheroid) from the donors 1 and 2 after different differentiation time points (2, 3 or 4 weeks) and from subcutaneous white adipose tissue (scWAT) and subcutaneous mature adipocyte fracture (MAF) from the donor 3, N = 1 per group. Data are presented as relative to scWAT = 1. Dashed line represents *PPARγ* expression level in MAFs.

For the donor characteristics, see Table 1. Note that in contrast to our gold standard protocol, adipocyte spheroids analysed in these experiments were not embedded into growth factor reduced-Matrigel.

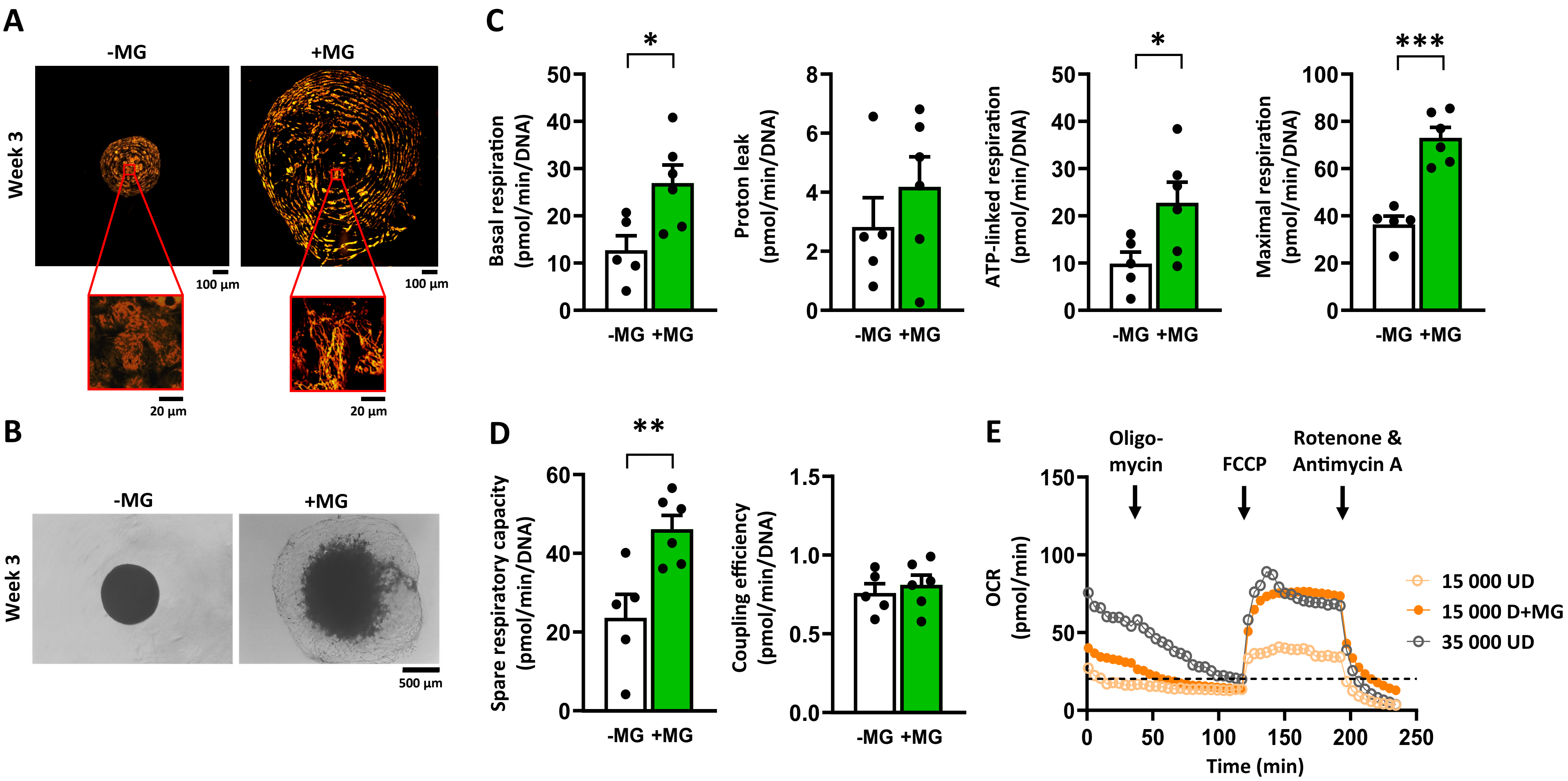

A
-MG
+MG
Week 3
100 µm
100 µm
20 µm
20 µm
B
-MG
+MG
Week 3
500 µm
C
Basal respiration (pmol/min/DNA)
Proton leak (pmol/min/DNA)
ATP-linked respiration (pmol/min/DNA)
Maximal respiration (pmol/min/DNA)
-MG +MG
*
*
***
D
Spare respiratory capacity (pmol/min/DNA)
Coupling efficiency (pmol/min/DNA)
**
-MG +MG
E
Oligo-mycin
FCCP
Rotenone & Antimycin A
OCR (pmol/min)
Time (min)
15 000 UD
15 000 D+MG
35 000 UD

**Supplementary Figure 2**. **Embedding of human subcutaneous white adipocyte spheroids into growth factor reduced (GFR)-Matrigel improves mitochondrial network and respiration.** A) Selected representative fluorescence live cell imaging of differentiated adipocyte spheroids derived from the donor 2 for the analysis of mitochondria (MitoTracker, yellow). On the left, adipocyte spheroid without (-MG) and on the right with (+MG) GFR-Matrigel embedding. Magnification 40x water immersion objective. B) Selected representative bright-field imaging on the differentiated adipocyte spheroids without and with GFR-Matrigel embedding. Parameters of mitochondrial respiration, including C) basal respiration, proton leak, ATP-linked respiration, maximal respiration and D) spare respiratory capacity and coupling efficiency in differentiated adipocyte spheroids without or with GFR-Matrigel embedding, N = 5-6 per group. E) Raw oxygen consumption rate (OCR) values from undifferentiated (UD) and/or differentiated embedded in Matrigel (D) (pre)adipocyte spheroids containing 15 000 or 35 000 cells per spheroid from the donor 2, N = 8-9 per group. For clarity, error bars are not shown. The dashed line represents the 20 pmol/min OCR level.

Data are shown as means with individual values. In C-D), unpaired t-test was used. * $P < 0.05$, ** $P < 0.01$ and *** $P < 0.001$.

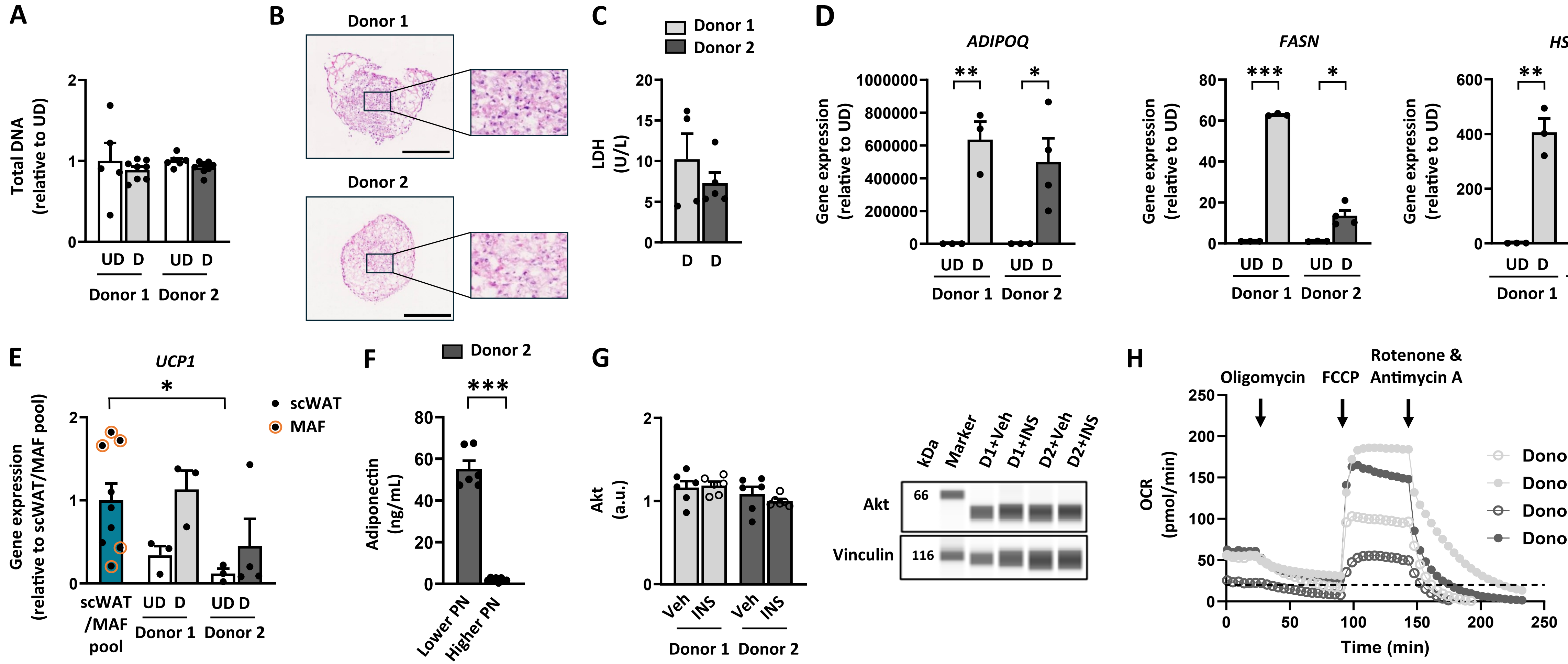

A
Total DNA (relative to UD)
UD D
Donor 1 Donor 2
B
Donor 1
Donor 2
C
Donor 1
Donor 2
LDH (U/L)
D
ADIPOQ
FASN
HSL
Gene expression (relative to UD)
UD D
Donor 1 Donor 2
E
UCP1
scWAT
MAF
Gene expression (relative to scWAT/MAF pool)
scWAT /MAF pool
Donor 1 Donor 2
F
Donor 2
Adiponectin (ng/mL)
Lower PN
Higher PN
G
Akt (a.u.)
Veh INS
Donor 1 Donor 2
kDa
Marker
D1+Veh
D1+INS
D2+Veh
D2+INS
Akt
66
Vinculin
116
H
Oligomycin
FCCP
Rotenone & Antimycin A
OCR (pmol/min)
Time (min)
Donor 1 UD
Donor 1 D
Donor 2 UD
Donor 2 D

**Supplementary Figure 3**. **Characteristics of undifferentiated (UD) and differentiated (D) (pre)adipocyte spheroids**. A) Total DNA amount in UD and D (pre)adipocyte spheroids derived from the donors 1 and 2, N = 5-8 per group. B) Selected representative cryosections with haemotoxylin and eosin staining images of the differentiated adipocyte spheroids from the donors 1 and 2. Scale bar = 500 µm. C) Lactate dehydrogenase (LDH) enzyme activity measured from the media of differentiated adipocyte spheroids derived from the donors 1 and 2 after the 3-week differentiation period, N = 4-5 per group. Gene expression of D) *ADIPOQ, FASN* and *HSL* in UD and D (pre)adipocyte spheroids derived from the donors 1 and 2, N = 3-4 per group. Data are presented as relative to UD = 1. Gene expression of E) *UCP1* in UD and D (pre)adipocyte spheroids relative to a pool of subcutaneous human white adipocyte tissue (scWAT) and subcutaneous human mature adipocyte fraction (MAF) samples, N = 3-9 per group. Note that scWAT/MAF pool consists of samples derived from the donors 4-9 (Table 1). Data are presented as relative to scWAT/MAF pool = 1. F) Media adiponectin concentration from differentiated adipocyte spheroids derived from the donor 2 with lower (2-3) and higher (6) passage number (PN), N = 6-8 per group. G) Protein content of total Akt in response to a 30-min vehicle (Veh, 1x PBS) or 100 nM insulin (INS) administration to differentiated adipocyte spheroids derived from the donors 1 (D1) or 2 (D2), N = 6 per group. Representative blots are shown on the right. Vinculin was used for data normalization. F) Raw oxygen consumption rate (OCR) values from UD and D (pre)adipocyte spheroids from the donors 1 and 2, N = 13-32 per group. For clarity, error bars are not shown. For the normalized mitochondrial respiration data, see Fig. 2A-B. The dashed line represents the 20 pmol/min OCR level.

Data are shown as means with individual values. In A), C-D) and F), unpaired t-test or Mann-Whitney U-test was used. In E and G), one-way ANOVA followed by Uncorrected Fisher's LSD or with Kruskal–Wallis test followed by Uncorrected Dunn's test was used. * $P < 0.05$ and *** $P < 0.001$.

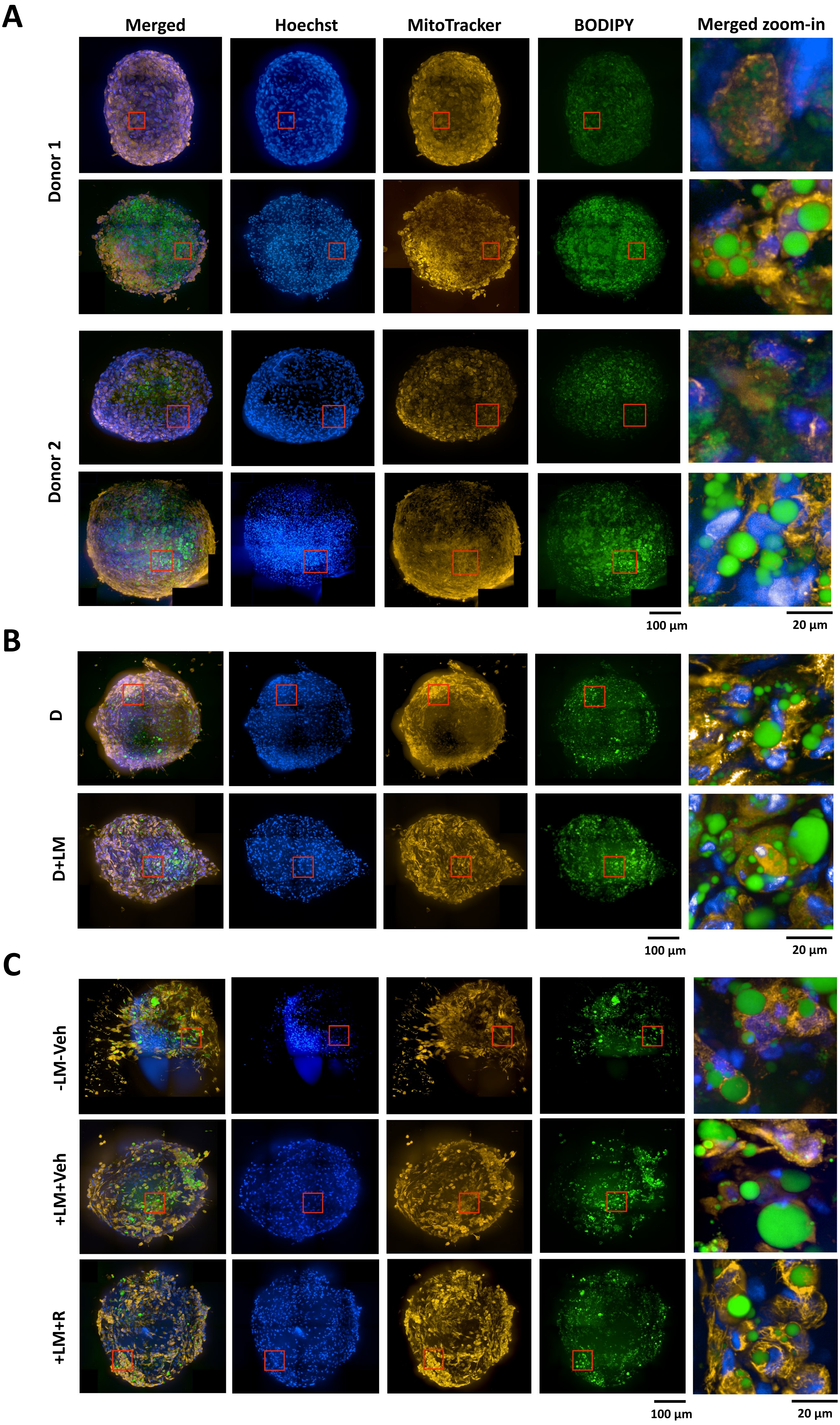

A
Merged
Hoechst
MitoTracker
BODIPY
Merged zoom-in
Donor 1
UD
D
Donor 2
UD
D
100 µm
20 µm
B
D
D+LM
100 µm
20 µm
C
-LM-Veh
+LM+Veh
+LM+R
100 µm
20 µm

**Supplementary Figure 4**. Selected representative fluorescence live cell imaging of the nuclei (Hoechst, blue), mitochondria (MitoTracker, yellow) and lipid droplets (BODIPY, green) and all three merged (whole adipocyte spheroid and zoom-in) in A) undifferentiated (UD) and differentiated (D) (pre)adipocyte spheroids derived from the donors 1 and 2, B) differentiated adipocyte spheroids administered without (D) and with lipid mixture (D+LM) derived from the donor 2 and C) differentiated adipocyte spheroids derived from the donor 2 administered without LM but with vehicle (-LM+Veh), with LM and vehicle (+LM+Veh) or with LM and 1 μM rosiglitazone (+LM+R) for 72h post differentiation. In C), DMSO was used as vehicle. In the figures, the red rectangle represents the area from which the merged zoom-in has been taken. Whole spheroid images; scale bar 100μm and merged zoom-ins; scale bar 20μm. Magnification 40x air objective.

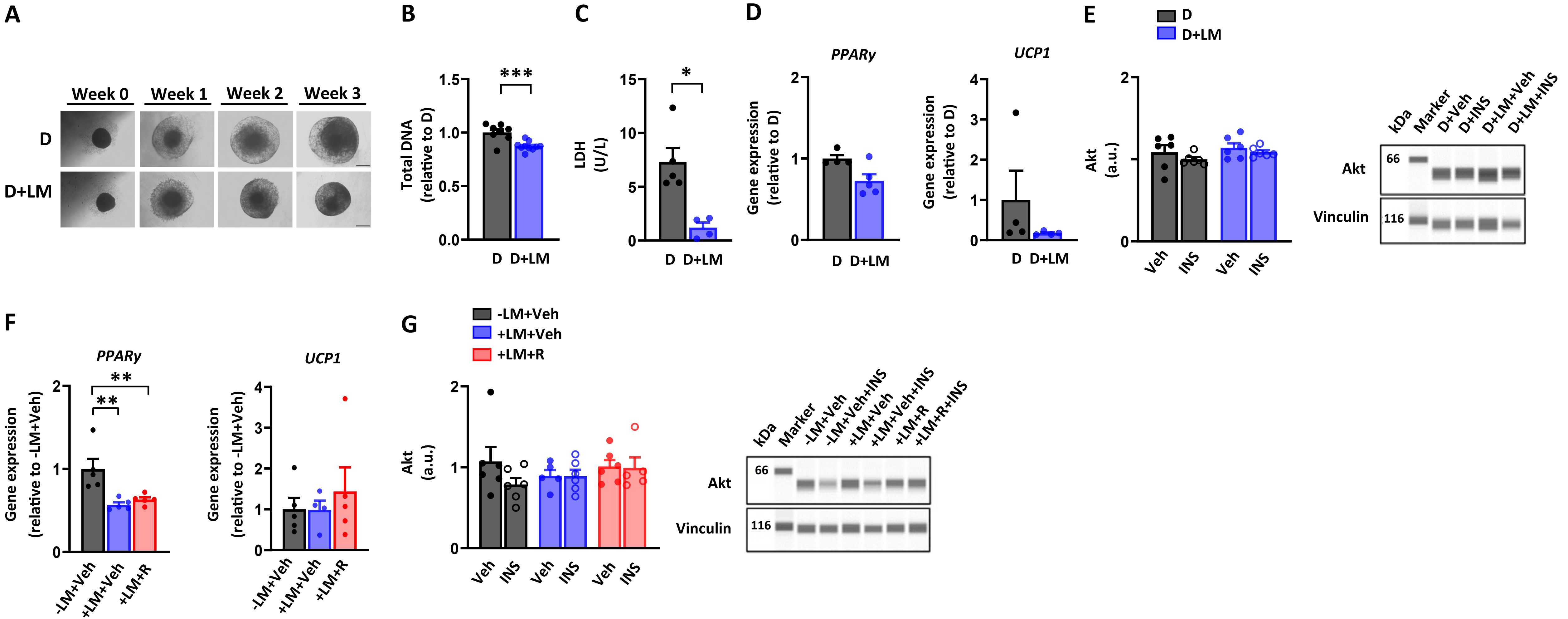

A
Week 0
Week 1
Week 2
Week 3
D
D+LM
B
Total DNA (relative to D)
***
D
D+LM
C
LDH (U/L)
*
D
D+LM
D
PPARγ
UCP1
Gene expression (relative to D)
D
D+LM
E
D
D+LM
Akt (a.u.)
Veh
INS
Veh
INS
kDa
Marker
D+Veh
D+INS
D+LM+Veh
D+LM+INS
Akt
Vinculin
66
116
F
PPARγ
UCP1
Gene expression (relative to -LM+Veh)
**
**
-LM+Veh
+LM+Veh
+LM+R
G
-LM+Veh
+LM+Veh
+LM+R
Akt (a.u.)
Veh
INS
Veh
INS
Veh
INS
kDa
Marker
-LM+Veh
-LM+Veh+INS
+LM+Veh
+LM+Veh+INS
+LM+R
+LM+R+INS
Akt
Vinculin
66
116

**Supplementary Figure 5**. **The effect of lipid mixture and/or rosiglitazone treatment on DNA content, cell viability and total Akt protein content.** Only differentiated adipocyte spheroids derived from the donor 2 were used in these experiments. A) Selected brightfield images of the adipocyte spheroids without (D) and with lipid mixture (LM) administration (D+LM) during the 3-week differentiation period. Scale bar = 500 µm. B) Total DNA amount in D and D+LM adipocyte spheroids, N = 8-10 per group. C) Lactate dehydrogenase (LDH) enzyme activity measured from D and D+LM adipocyte spheroid media after the 3-week differentiation period, N = 4-5 per group. Gene expression of D) *PPARγ* and *UCP1* in D and D+LM adipocyte spheroids, N = 3-4 per group. E) Protein content of total Akt in response to a 30-min vehicle (Veh, 1x PBS) or 100 nM insulin (INS) administration, N = 6 per group. Representative blots are shown on the right. Gene expression of F) *PPARγ* and *UCP1* in differentiated adipocyte spheroids administered with vehicle (-LM+Veh), with LM and vehicle (+LM+Veh) or with LM and 1 µM rosiglitazone (+LM+R), N = 4-5 per group. G) Protein content of total Akt in response to a 30-min vehicle (Veh, 1x PBS) or 100 nM insulin (INS) administration, N = 5-6 per group. Representative blots are shown on the right. In E and G), Vinculin was used for data normalization. In B) and D), data are presented as D = 1 and in F), -LM+Veh = 1.

Data are shown as means with individual values. In B-D), unpaired t-test or Mann-Whitney U-test was used. In E-G), one-way ANOVA followed by Uncorrected Fisher's LSD or with Kruskal–Wallis followed by Uncorrected Dunn's test was used. In figures, * $P < 0.05$, ** $P < 0.01$ and *** $P < 0.001$.
